# Longitudinal Multi-Organ Transcriptomic Atlas of Salt-Induced Hypertension

**DOI:** 10.1101/2025.08.27.672714

**Authors:** Ratnakar Tiwari, Olha Kravtsova, Lashodya V. Dissanayake, Melissa Lowe, Biyang Xu, Vladislav Levchenko, Steven Didik, Ruslan Bohovyk, Daria V. Ilatovskaya, Oleg Palygin, Alexander Staruschenko

## Abstract

**BACKGROUND:** Salt-sensitive hypertension is a prevalent and clinically significant subtype of hypertension, where increased dietary salt intake elevates blood pressure and causes injury to multiple organ systems. Despite extensive research, dynamic molecular changes and conserved versus organ-specific transcriptional programs in hypertensive multi-organ damage remain poorly understood. Defining complex molecular pathways both in a temporal sequence and in an organ-specific manner is essential for developing targeted, precision therapies to mitigate hypertensive disease burden.

**METHODS:** We generated a longitudinal multi-organ transcriptomic atlas of salt-sensitive hypertension using RNA sequencing of kidney cortex, kidney medulla, heart, and liver from Dahl salt-sensitive rats across four disease stages. A comprehensive bioinformatic analysis mapped dynamic transcriptional programs, evaluated 50 biological pathways, and defined upstream regulators. Histological and biochemical assays complemented transcriptomic analysis, while integration with human genome-wide association studies (GWAS) and compound–transcriptome analysis provided translational insights and identified candidate therapeutics.

**RESULTS:** Salt-induced hypertension elicited both shared and tissue-specific transcriptional programs that evolved with disease progression. The kidney medulla showed robust early immune activation with metabolic suppression, while the cortex exhibited transient metabolic activation before declining and initiating immune activation. The liver and heart showed time-dependent metabolic and inflammatory remodeling. Cross-organ comparisons revealed a shared early proliferative response that converged on proinflammatory and fibrotic signatures. Upstream regulator analysis identified 79 time- and tissue-specific transcription factors associated with gene expression dynamics. GWAS integration analysis revealed endocrine signaling, ion transport, lipid metabolism, and detoxification as conserved pathways across species, underscoring the translational relevance of the model and study. Predictive compound– transcriptome analyses identified kinase inhibitors targeting phosphoinositide 3-kinase, mechanistic target of rapamycin and cyclin-dependent kinases as top candidates to counteract maladaptive transcriptional programs.

**CONCLUSIONS:** This study defines temporal and tissue-specific transcriptomic remodeling in salt-sensitive hypertension and highlights the need for precision interventions to prevent progressive organ damage.

## INTRODUCTION

Hypertension is one of the most prevalent and major cardiovascular risk factors globally. It affects over one billion people and significantly contributes to stroke, heart failure, and chronic kidney disease (CKD)^1–4^. Among its subtypes, salt-sensitive hypertension (SSH) affects up to 30% of the global adult population and more than half of all individuals with hypertension and is strongly associated with CKD^5–7^. Further, this burden disproportionately impacts high-risk groups, including African Americans, individuals with metabolic syndrome, and patients with pre-existing renal disease^8^. Despite its prevalence and clinical impact, SSH remains mechanistically complex and difficult to diagnose and manage^7,9–11^.

Although SSH has traditionally been studied from a renal-centric perspective, its impact extends beyond the kidney and cardiovascular system. Among other affects organs, the liver is increasingly recognized as a target of SSH-induced metabolic reprogramming, with hepatic gene expression changes that contribute to cardiovascular disease (CVD)^12,13^. While these studies have significantly advanced our understanding of how SSH affects specific organs, they primarily focus on individual organs at selected time points. As a result, they fall short of capturing the full complexity of SSH as a systemic and dynamically evolving disease. This underscores a pressing need for longitudal, multi-organ studies that can illuminate the cross-organ as well as organ-specific molecular trajectories that drive disease progression over time across multiple organs.

To address this critical knowledge gap, we conducted a comprehensive, temporally resolved, multi-organ transcriptomic atlas of SSH in the Dahl salt-sensitive (SS) rat model. This model remains one of the most physiologically relevant and widely utilized preclinical models of SSH, recapitulating key human features such as progressive hypertension, renal injury, and maladaptive cardiovascular remodeling in response to a high salt diet^14–17^. Previous work from our group and others leveraging different cellular, molecular, and omics approaches in this model has uncovered several important aspects of disease progression, including the activation of immune signaling pathways and metabolic reprogramming primarily within the kidney at specific stages of SSH^18–22^. However, the temporal dynmics of these processes across organs has not been systematically characterized.

Using advanced bioinformatics and network-based modeling, we mapped dynamic activation and suppression of molecular pathways underlying SSH across multiple tissues. This revealed both conserved and tissue-specific molecular pathways, as well as central hub regulators that drive maladaptive responses against sustained SSH. Integration with genome-wide association studies (GWAS) and druggable pathway prioritization highlighted actionable therapeutic targets with potential for drug repurposing and precision intervention. Integrated with histological and biochemical analyses, this longitudinal multi-organ transcriptomic atlas delivers a systems-level view of gene expression changes in SSH and serves as a resource for understanding tissue- and time-specific mechanisms that contribute to chronic end-organ damage.

## METHODS

Detailed methods are provided in the supplemental materials.

### Data availability

Data can be explored on our online data resource at *STAR LAB-Hypertension Atlas* (https://starlabusf.com/). Additional data and supporting materials are available from the corresponding authors upon reasonable request.

## RESULTS

### A longitudinal transcriptomic atlas of salt-sensitive hypertension

To systematically define the transcriptional landscape underlying SSH, we performed mRNA sequencing across four tissues: kidney cortex, kidney medulla, liver, and heart. Samples were collected from Dahl SS rats fed a high salt (HS, 4% NaCl, # D113756, Dyets Inc) diet at four progressive stages of hypertension (days 7, 14, 21, and 35), alongside a normal salt diet control (NS, 0.4% NaCl, # D113755, Dyets Inc). This design spans the full course of disease progression, beginning with the onset of high blood pressure by day 7 (Figure S1A), through to late-stage pathology, including the terminal phase at day 35 when animals start to show advanced cardiovascular dysfunction leading to stroke and death. In total, the study yields 120 high-quality transcriptomes, providing a rich resource to decode the longitudinal multi-organ molecular dynamics of SSH (Figure 1A).

**Figure 1.**
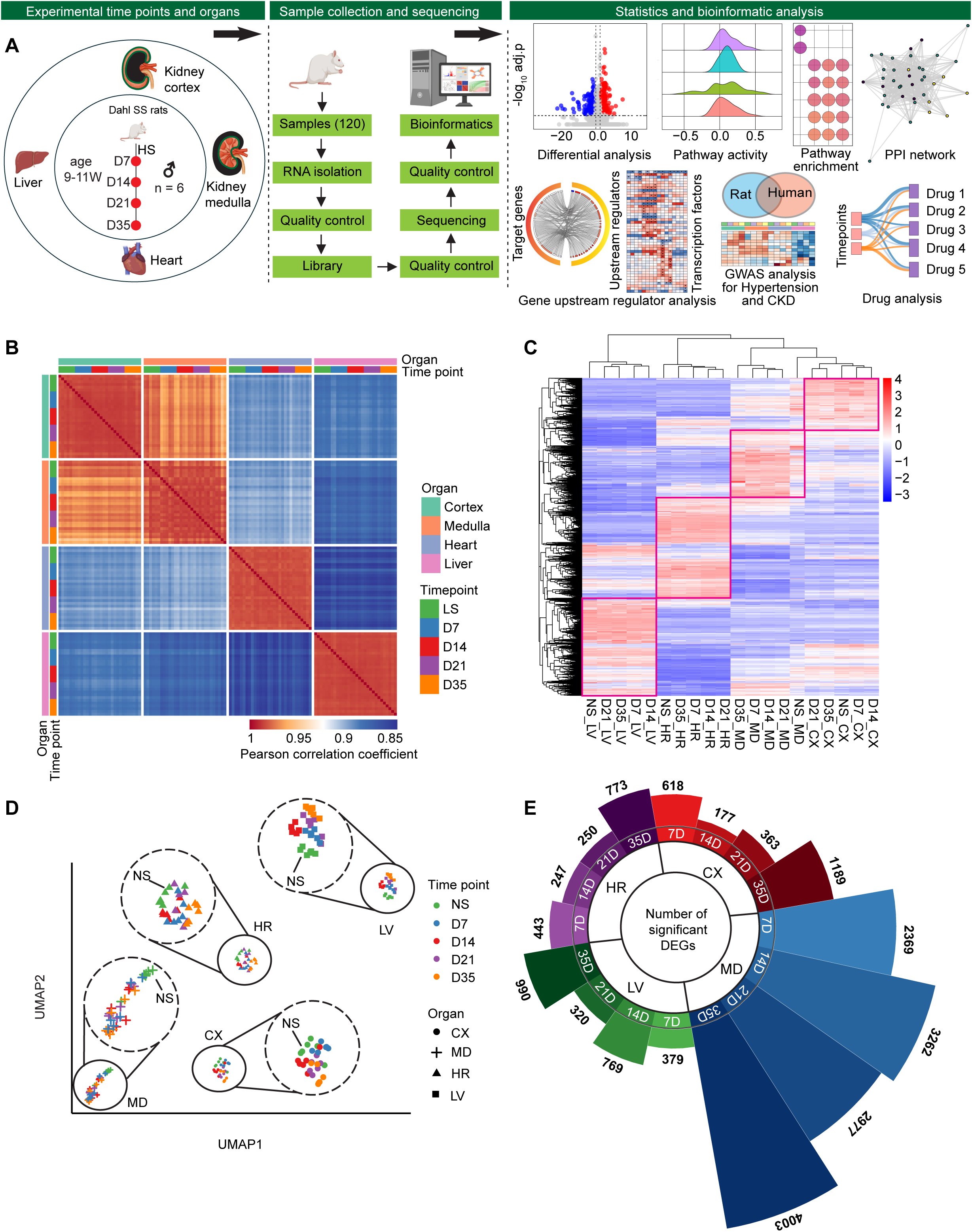
Overview of experimental design, data quality, and global transcriptional profiling. **A**. Schematic of the study design and bioinformatics pipeline. Male Dahl SS rats were maintained on a normal salt (NS) control diet or switched to a high salt (HS) diet at 9–11 weeks of age to induce hypertension and organ injury. Kidney cortex, kidney medulla, liver, and heart samples were collected from NS controls and HS-fed rats after 7, 14, 21, and 35 days of treatment. RNA was isolated, quality-checked, and sequenced. Reads underwent sample-level quality control, alignment, and quantification. Downstream analyses included variance-stabilizing transformation (VST), differential expression analysis, pathway and upstream regulator enrichment, protein–protein interaction network analysis, genome-wide association study (GWAS) integration, and drug target analyses. **B**. Sample–sample Pearson correlation heatmap, annotated by organ and time point. **C**. Heatmap of all significantly differentially expressed genes (adjusted *P* < 0.05), hierarchically clustered by sample (columns) and gene (rows). Red boxes highlight major co-expression gene modules segregating primarily by organ. **D**. UMAP embedding of all 120 samples using VST counts, showing predominant clustering by organ. Dashed circles highlight zoomed-in views of distinct tissue clusters, illustrating separation of NS from HS experimental groups. **E**. Radial bar chart summarizing the number of differentially expressed genes (|log₂ fold change| ≥ 0.585, equivalent to a fold change ≥ 1.5, adjusted *P* < 0.05) in each tissue and time point, illustrating dynamic and tissue-specific transcriptional responses to HS-induced hypertension. CX, cortex; MD, medulla; LV, liver; HR, heart; D7, day 7; D14, day 14; D21, day 21; and D35, day 35 time points. *n* = 6 male rats per group.

Principal component analysis (PCA) of variance-stabilized gene expression profiles revealed strong segregation by organ identity. The first two components explained over 80 percent of the total variance (Figure S1B). Kidney cortex and medulla samples formed adjacent but distinct clusters, while liver and heart resolved along orthogonal axes, reflecting global transcriptomic variation. Rank-abundance plots confirm stable expression distributions across samples and time points (Figure S1C), with no apparent loss of complexity or RNA integrity, further affirming data quality. Sample-sample Pearson correlation matrices also revealed hierarchical structure dominated by organ identity (Figure 1B). These correlations remained consistently high (Pearson correlation coefficient r > 0.95 within organs), underscoring excellent reproducibility and biological coherence. Among organs, kidney cortex and medulla were the most closely related (r > 0.98), while more moderate correlations were observed between renal and non-renal tissues (Data sheet S1). Hierarchical clustering of all genes (adjusted *P* < 0.05), based on log₂-transformed expression levels, also partitioned samples into well-defined blocks corresponding to organ (Figure 1C). Uniform manifold approximation and projection (UMAP) was also applied to the expression matrix to visualize global variance and local structure in transcriptional profiles (Figure 1D). In UMAP analysis, distinct organ-specific clusters also showed separation between non-hypertensive conditions (control, NS) to different stages of SSH remodeling.

Next, we performed differential expression analysis for each time point, comparing tissue samples to corresponding NS control tissue. Applying a threshold of adjusted *P* value less than 0.05 and absolute log₂ fold change greater than or equal to 0.585 (corresponding to absolute fold change ≥ 1.5), we identified a significant number of differentially expressed genes (DEGs) across the datasets (Figure S2). We found a dynamic transcriptional pattern with an early elevation in DEGs, a mid-stage decline, and a subsequent rise at the later stage of SSH (Figure 1E). Notably, the kidney medulla exhibited greater magnitude changes than the other tissues, with the number of DEGs increasing sharply from 2,369 on day 7 to 3,262 on day 14, decreasing to 2,977 on day 21, and peaking at 4,003 by day 35. A similar pattern was detected for the liver. However, for the kidney cortex and heart, the number of DEGs showed transient suppression of initial response earlier, on day 14, with following reactivation also peaking by day 35. These phased trajectories suggest differential responses across tissues, with the kidney medulla exhibiting the highest vulnerability.

To better understand the physiological context of these transcriptional changes, we next examined systemic electrolyte balance. The high salt diet group showed a marked increase in sodium and chloride excretion (Figure S3A–B). However, blood sodium levels remained normal across all time points, whereas blood chloride levels exhibited a consistent reduction from day 7 onwards (Figure S3C–D). These changes were accompanied by a significant increases in blood pH, indicating a systemic change in acid–base homeostasis (Figure S3E). To see how transcriptomic changes relate to organ damage, we looked at organ weights and observed an increase in relative kidney weight beginning at day 7. Further, we observed a marked diuresis and a progressive rise in urinary albumin excretion, which peaked at day 21 before partially declining by day 35 (Figure 2A–C). Notably, serum creatinine levels became only significantly elevated at the late stage, on day 35 (Figure 2D). Next, we performed comprehensive histopathological assessments of the kidney, liver, and heart across all time points. The medulla exhibited severe and progressive cellular damage, cast formation, and fibrosis. In contrast, the kidney cortex showed moderate injury, with lower levels of cast formation and fibrosis (Figure 2E–K). The changes in kidney were consistent with the expression of genes associated with kidney injury, inflammation, and fibrosis (Figure 2L). Histopathological analysis of the liver and heart revealed perivascular fibrosis in Masson’s trichrome staining, consistent with the notable changes in the inflammatory and fibrosis-associated genes (Figure S3F–H).

**Figure 2.**
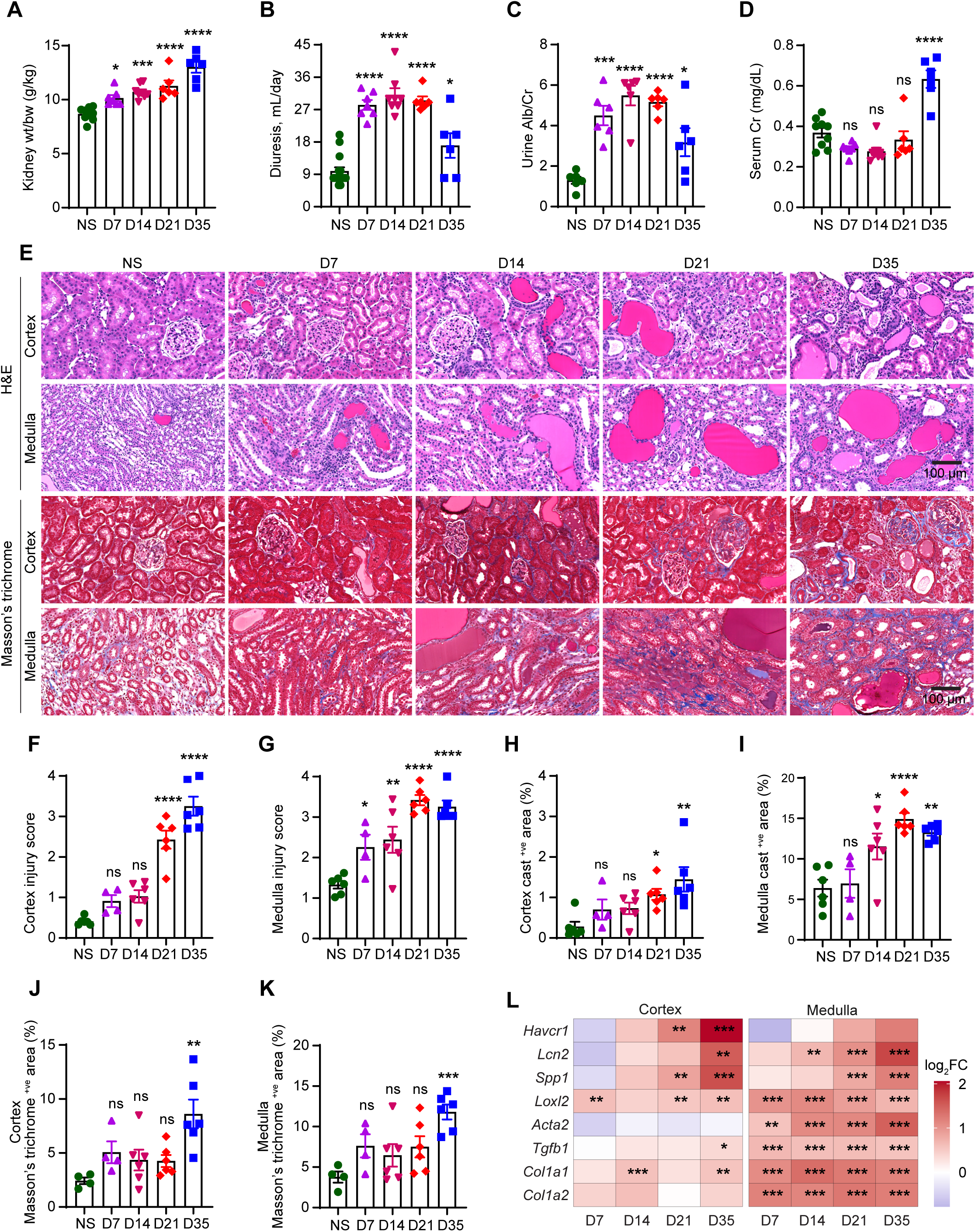
High salt diet-induced hypertension promotes progressive, region-specific injury in the kidney. Dahl SS rats were maintained on a normal salt diet (NS; 0.4% NaCl) diet or switched to a high salt (HS; 4% NaCl) diet for 7, 14, 21, or 35 days. **A**. Kidney weight normalized to body weight (g/kg). **B**. 24-hour urine volume (mL/day). **C**. Urine albumin-to-creatinine ratio (Alb/Cr). **D**. Serum creatinine concentration (mg/dL). **E**. Representative hematoxylin and eosin (H&E; top two rows) and Masson’s trichrome (bottom two rows) staining of renal cortex and medulla from NS- and HS-fed rats at the indicated time points (scale bars show 100 µm). **F–G**. Injury scores in cortex (**F**) and medulla (**G**). **H–I**. Percentage of cast-positive area in cortex (**H**) and medulla (**I**). **J–K**. Percentages of Masson’s trichrome positive area in cortex (**J**) and medulla (**K**). Data are shown as mean ± standard error of the mean (SEM); *n* = 4–15 male rats per group. For **A–D** and **F–K**, one-way ANOVA with Šídák’s multiple comparisons was used. \**P* < 0.05; *\*\*P* < 0.01; \*\*\**P* < 0.001; \*\*\*\**P* < 0.0001; ns, not significant. **L**. Heatmaps of selected genes associated with injury, fibrosis and inflammation showing log₂ fold-change (vs NS) at each time point in cortex and medulla. Expression of genes was extracted from RNA-seq analysis. On the heatmap, asterisks denote Benjamini–Hochberg adjusted *P* thresholds (*adj. P*). \**adj. P* < 0.05; *\*\*adj. P < 0.01;* \*\*\**adj. P* < 0.001; *n* = 6 male rats per group. D7, day 7; D14, day 14; D21, day 21; and D35, day 35 time points. *Havcr1*, *hepatitis A virus cellular receptor 1/kidney injury molecule-1*; *Lcn2*, *lipocalin-2*; *Spp1*, *secreted phosphoprotein 1*; *Loxl2, lysyl oxidase-like 2*; *Acta2, actin alpha-2*, *smooth muscle*; *Tgfb1*, *transforming growth factor beta 1*; *Col1a1, collagen type I alpha-1 chain*; *Col1a2*, *collagen type I alpha-2 chain*.

Collectively, this longitudinal multi-organ transcriptomic atlas, supported by biochemical assays and histopathology, defines the dynamic transcriptomic, cellular, and functional landscape of SSH. It captures both early and long-term changes in gene expression across the kidney, liver, and heart.

### Salt-induced hypertension drives dynamic and tissue-specific reprogramming of core biological pathways

To identify coordinated molecular responses to SSH, we quantified the activity of 50 Hallmark pathways in different tissues over time. This analysis revealed coordinated activation and repression of canonical pathways with disease progression, uncovering distinct longitudinal multi-organ pathway activity patterns (Figure 3A, Figure S4). As aforementioned kidney medulla exhibited the most robust and sustained changes in gene diversity and pathway activity. By day 7, inflammatory pathways, including cytokine signaling IL6–JAK–STAT3 pathway, interferon alpha, and interferon gamma were strongly upregulated and remained elevated across all subsequent time points. In parallel, proliferative and stress-adaptive programs, such as E2F targets, G2M checkpoint, and mitotic spindle pathways, were also activated early and sustained. In contrast, metabolic pathways, including oxidative phosphorylation and fatty acid metabolism, were markedly suppressed at day 7 and remained inhibited as compared to the NS control medulla group. Developmental programs such as angiogenesis and epithelial–mesenchymal transition (EMT) were persistently elevated, suggesting continuous tissue remodeling responses. However, the kidney cortex showed comparibly moderate response. Metabolic programs such as oxidative phosphorylation were slightly induced on day 7, followed by a decline over time. Inflammatory pathways, including complement, IL6–JAK–STAT3, and allograft rejection, were gradually upregulated, reaching peak activity at later time points. Stress pathways such as hypoxia followed a similar pattern, with late-stage elevation. Structural remodeling pathways (angiogenesis, EMT) also followed a delayed activation trajectory.

**Figure 3.**
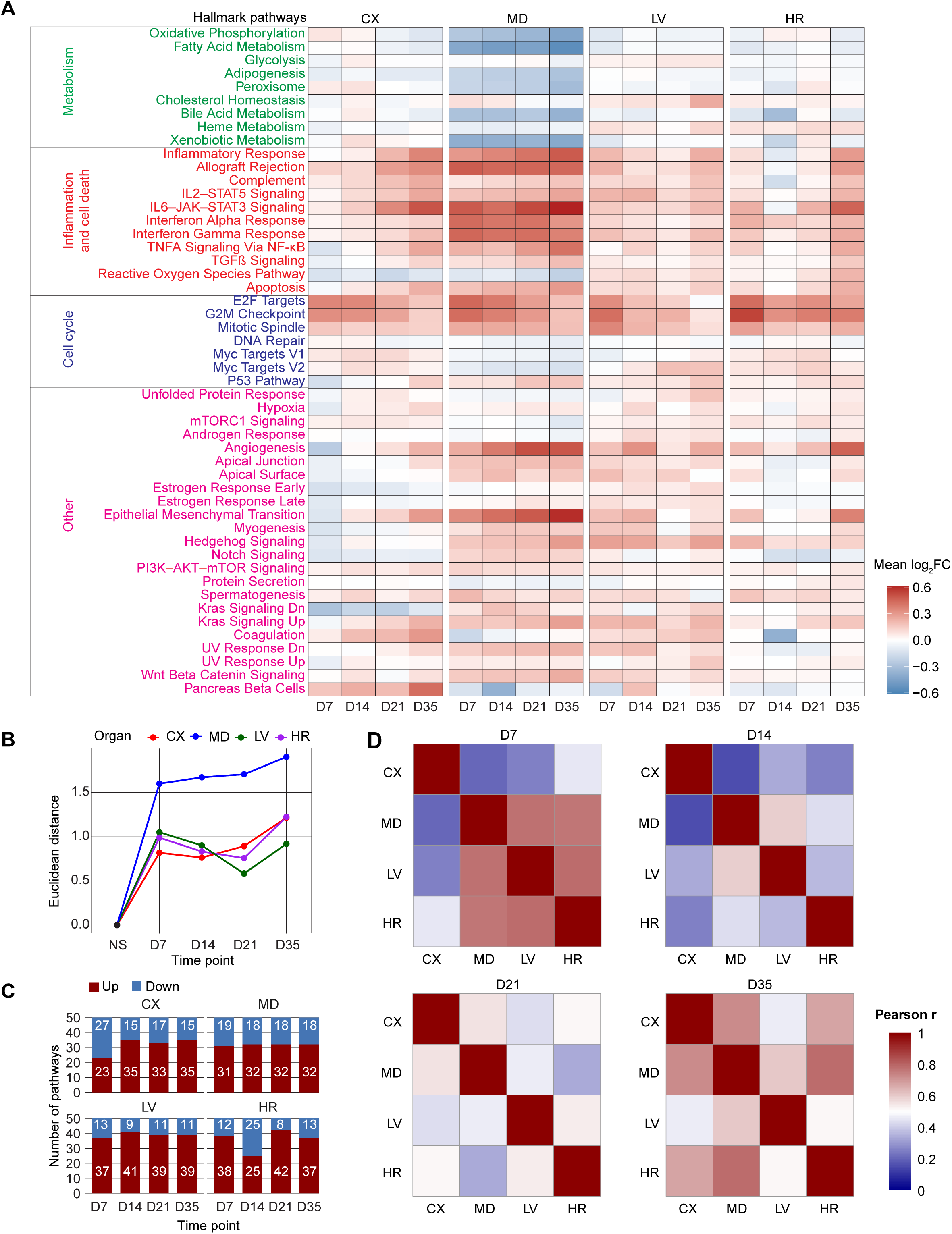
High salt diet-induced hypertension dynamically regulates biological pathway activity across organs. **A**. Mean pathway activity heatmap of 50 Hallmark pathways across cortex, medulla, heart, and liver at days 7, 14, 21, and 35 after initiation of a high salt (HS) diet compared to normal salt (NS) controls. Pathways are highlighted as functional groups, including metabolism, inflammation and death, cell cycle, and other clustered along the y-axis. **B**. Euclidean distances from baseline pathway profiles plotted over time, representing the magnitude of transcriptomic remodeling in each organ. The medulla shows the greatest divergence, followed by the heart, cortex, and liver. **C**. Stacked bar charts enumerating up-regulated (red) and down-regulated (blue) Hallmark pathways at each time point for each organ. Numeric labels indicate pathway counts, highlighting distinct kinetics of pathway engagement across tissues. **D**. Heatmaps showing Pearson correlation between pathway profiles across tissues (cortex, medulla, heart, and liver) at each time point (days 7, 14, 21, and 35). Each panel compares all organ pairs for a single time point, revealing similarities and differences in pathway responses among organs after the HS diet. CX, cortex; MD, medulla; LV, liver; HR, heart; D7, day 7; D14, day 14; D21, day 21; and D35, day 35 time points. *n* = 6 male rats per group.

In the liver, several metabolic programs, including glycolysis, cholesterol homeostasis, and heme metabolism, together with proliferative pathways such as E2F targets and the G2M checkpoint, were upregulated as early as day 7 and remained elevated throughout the study period. Immune and inflammatory pathways also showed persistent activation, although at a lower magnitude than in the kidney. The heart exhibited modest activation of selected immune and stress-related pathways at day 7, followed by a transient attenuation at day 14. By day 21, a second wave of pathway activity emerged, marked by the induction of inflammatory-related pathways (allograft rejection, IL2–STAT5, IL6–JAK– STAT3), metabolic programs (cholesterol homeostasis), and structural remodeling signals (angiogenesis and EMT).

To quantify how far each tissue’s global pathway profile deviated from its baseline state, we calculated Euclidean distances (Figure 3B). This analysis measures the overall magnitude of change across all pathways relative to control. It confirmed the medulla’s early and sustained divergence, while the cortex, liver, and heart followed more gradual trajectories with moderate changes. To assess the directionality of biological pathway regulation over time, we quantified the number of upregulated and downregulated Hallmark pathways in each tissue (Figure 3C). This analysis revealed that the renal cortex exhibited a progressive shift toward increased pathway activation, with an early rise in up-regulated pathways accompanied by a sustained decline in down-regulated pathways. The medulla showed a consistent pattern over time, with relatively stable numbers of both up- and down-regulated pathways. The liver demonstrated predominantly up-regulated pathways with minimal evidence of repression, indicating sustained transcriptional activation. In contrast, the heart displayed a dynamic response, characterized by early activation, a transient reduction in up-regulated pathways at intermediate time points, and renewed activation at later stages. These findings further highlight distinct, tissue-specific kinetics of pathway-level reprogramming during the progression of SSH.

To better understand whether different organs follow similar or distinct patterns of biological pathway activities during the progression of SSH, we calculated Pearson correlations using activity scores of 50 Hallmark pathways at each time point (Figure 3B, Data sheet S2). Notably, till day 14, the kidney medulla and cortex exhibited no significant correlation in pathway activity (D7: r^2^ = 0.04, *P* = 0.14; D14: r^2^ = 0.02, *P* = 0.28), suggesting distinct molecular responses despite their anatomical proximity. Interestingly, the medulla showed some similarity in pathway responses on day 7 with extra-renal tissues, showing significant correlations observed with the liver (D7: r^2^ = 0.60, *P* <0.0001) and heart (r^2^ = 0.57, *P* <0.0001) which droped on day 14 (liver: r^2^ = 0.35, *P* <0.0001; heart: r^2^ = 0.18, *P* <0.01). On day 21, the medulla and cortex started showing similarities in pathway activity (r^2^ = 0.3, *P* <0.0001) while the pathway activity similarities between medulla with liver and heart droped (Data sheet S2). On day 35, the similarities in pathway activities increased again with medulla showing significant correlations with cortex (r^2^ = 0.53, *P* <0.0001), liver (r^2^ = 0.37, *P* <0.0001) and heart (r^2^ = 0.62, *P* <0.0001). Overall, this analysis indicates that in the early stages of SSH, tissues engage distinct biological pathway responses, with the medulla aligning more with extra-renal tissues than with the adjacent cortex. Over time, biological pathway dynamics converged across tissues, reflecting progression to a systemic state of chronic injury.

To delineate the organizational structure of transcriptional responses across organs and time, we applied UMAP dimensionality reduction to the Hallmark pathway activity matrix encompassing all tissue– time point combinations, followed by unsupervised clustering (Figure S5). This approach grouped pathways based on similarity in temporal dynamics, independent of absolute expression levels or tissue origin. By embedding pathway trajectories in a low-dimensional space, UMAP preserved relative regulatory relationships, enabling pathways with comparable dynamics across organs to cluster together. The resulting modules captured coherent biological themes, including mitochondrial and lipid metabolism, immune and inflammatory signaling, cell cycle progression, and stress adaptation. The distinct spatial separation and tight grouping of clusters reflected coordinated engagement of different biological programs during salt-induced hypertension, indicating synchronized activation of metabolic, inflammatory, and remodeling processes throughout disease progression.

In summary, the analysis of 50 hallmark pathways demonstrates that SSH induces distinct tissue-specific remodeling in the kidney, heart, and liver, while also engaging a subset of pathways that exhibit cross-organ involvement, suggesting partial convergence of molecular responses during disease progression.

### Shared molecular signatures uncover early cell cycle activation and late immune remodeling across organs

To uncover conserved transcriptional programs in SSH across tissues, we identified common genes that were significantly differentially expressed in all four tissues (cortex, medulla, liver, and heart) and can represent global targets at each time point of the disease progression. Venn diagrams revealed a dynamic change of organ-specific and shared responses (Figure 4A). The early transcriptional response at day 7 showed a set of 44 genes common to all tissues. This shared signature decreased at days 14 and 21, but a distinct set of 33 overlapping genes reappeared at day 35. Heatmaps of these common DEGs demonstrated consistent expression patterns across organs within each time point (Figure 4B, F–H). Analysis of the D7 common genes using Hallmark and Gene Ontology Biological Process (GO-BP) revealed a focused cell cycle program, including G2M checkpoint, E2F targets, and mitotic spindle terms (Figure 4C and Figure S6A and Data sheet S3). These were further supported by semantic similarity-based network analysis, which grouped top GO-BP terms into connected modules (Figure S6B).

**Figure 4.**
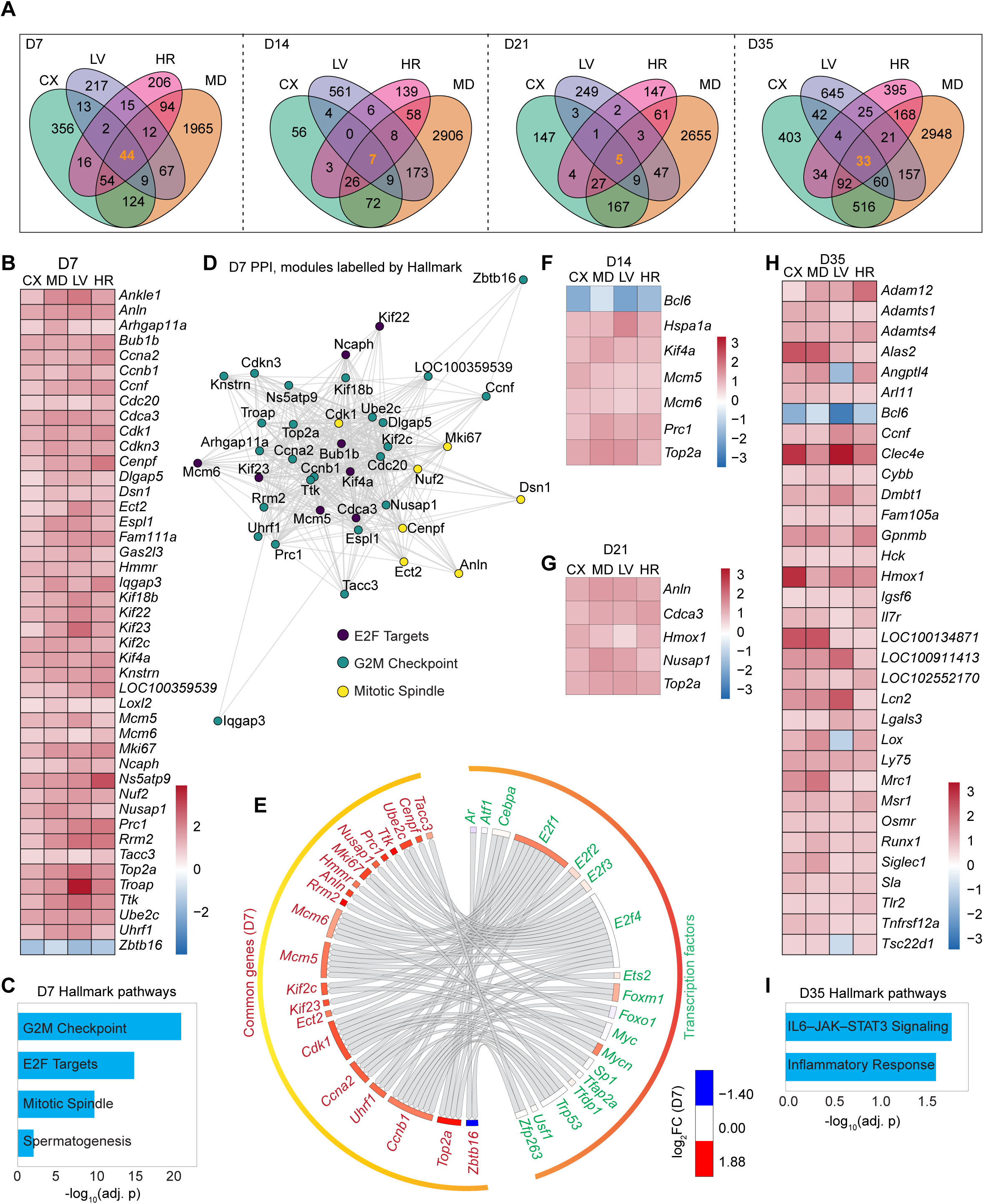
Shared transcriptional and pathway programs across organs in response to high salt diet-induced hypertension. **A**. Venn diagrams show overlap of DEGs across cortex, medulla, liver, and heart at each time point (days 7, 14, 21, and 35). Each segment displays the number of unique and shared DEGs. Genes shared by all tissues are highlighted in center. **B**. Heatmap of common 44 DEGs genes across all tissues on day 7. **C**. Bar plot showing significant Hallmark pathways emerged in the analysis of day 7 common genes. **D**. STRING-derived protein-protein interaction (PPI) network of common day 7 genes, with modules annotated by top enriched Hallmark terms per Louvain clustering. **E**. Circos plot showing high-confidence TF–target interactions for day 7 common genes, where outer sectors represent TFs and targets, colored by average log₂FC. Directional links denote regulatory interactions. **F–H**. Heatmaps of common DEGs at days 14, 21, and 35. **I**. Hallmark pathways emerged for D35 common genes. CX, cortex; MD, medulla; LV, liver; HR, heart; D7, day 7; D14, day 14; D21, day 21; and D35, day 35 time points. adj. p indicated adjusted *P* value. *n* = 6 male rats per group.

To examine the organizational architecture of identified global targets, we constructed protein– protein interaction (PPI) networks using STRING^23^. The D7 common genes formed a densely connected network, which resolved into distinct modules (Figure 4D), each enriched for canonical cell cycle processes. Notably, these modules were independently labeled by Hallmark terms via community-level overrepresentation analysis, reflecting their coordinating roles in cell division. To further dissect regulatory architecture, we integrated high-confidence transcription factor (TF)–target interactions from the DoRothEA regulon^24^ with day 7 common genes. This analysis revealed 18 TFs, including *forkhead box M1 (Foxm1*), *MYC proto-oncogene* (*Myc*), and *E2F transcription factor 3* (*E2f3*), which converged on 22 common differentially expressed genes (Figure 4E). Notably, several target genes, including *cyclin A2* (*Ccna2*), *DNA topoisomerase II alpha* (*Top2a*), and *cyclin-dependent kinase 1* (*Cdk1*), were co-regulated by multiple TFs, underscoring a core network of transcriptional convergence tightly linked to mitotic control. In contrast, the shared gene program on day 35 displayed a stark shift in composition and function. The 33 common genes included immunomodulators such as *toll-like receptor 2* (*Tlr2*) and *sialic acid binding Ig like lectin 1* (*Siglec1*), extracellular matrix regulators including *a disintegrin and metalloproteinase with thrombospondin motifs 1 (Adamts1*) and *lysyl oxidase (Lox*), and transcriptional repressors *B-cell lymphoma 6* (*Bcl6*) and *runt related transcription factor 1* (*Runx1*). The enrichment results at this time point indicated cytokine signaling IL6-JAK-STAT3 and vascular remodeling pathways (Figure 4I and Figure S6C). The GO-BP terms were more sparsely connected than on day 7, indicating a broader heterogeneity (Figure S6D and Data sheet S3). Consistently, the day 35 PPI network did not resolve into a well-connected structure (data not shown), marking a diversified immunometabolic signature.

In summary, we identified conserved, time-dependent transcriptional programs that define multi-organ remodeling in SSH. An early proliferative response was shared across organs, whereas later stages featured a more heterogeneous immune and matrix remodeling signature. These conserved molecular targets offer potential avenues for pharmacological strategies to mitigate multi-organ injury in SSH.

### Organ-unique transcriptional profiles reveal complexity of SSH

Step aside from the global transcriptomic signatures we next looked to more organ specific gene remodeling in the development of SSH. We identified organ-unique responder genes, defined as differentially expressed only in a single tissue across all time points compared to its respective control. This analysis revealed a robust, tissue-specific transcriptomic chnages (Figure 5A). The kidney medulla accounted for the largest number of organ-unique genes (3,645), followed by the liver (935), cortex (459), and heart (438), highlighting the medulla’s pronounced transcriptional responsiveness to salt-induced hypertension (Data sheet S4). Next, we defined genes that remained differentially expressed across all four time points (day 7–35) within a single tissue (stable unique genes). This subset contained 673 genes, further dominated by the medulla (644 genes), with only limited representation in liver (14), heart (11), and cortex (4) (Figure 5B and Data sheet S5). These genes represent persistent, tissue-specific responders to SSH.

**Figure 5.**
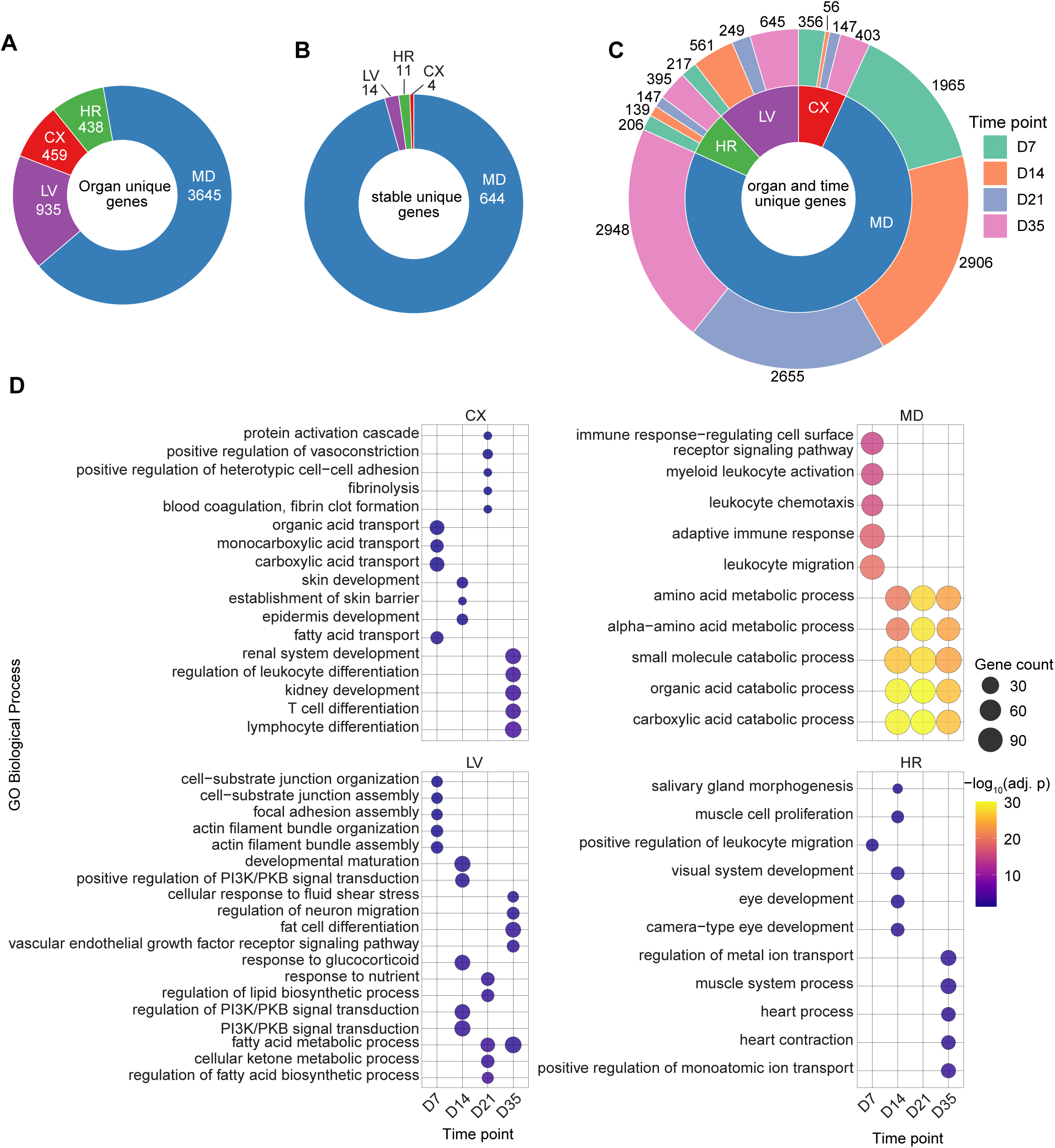
High salt diet-induced hypertension drives distinct temporal and organ-specific transcriptional programs, revealing tissue-specific injury mechanisms. **A**. Donut plot showing the number of DEGs that are unique to each tissue across any time point. **B**. Subset of tissue-specific genes that are consistently differentially expressed across all four time points within a given tissue. **C**. Nested donut chart illustrates the number of DEGs unique to each tissue and time point. Inner ring represents tissue, while the outer ring breaks them down by time point (days 7–35) and showing unique DEGs per tissue per time point. **D**. Top five significantly enriched GO Biological Process (GO-BP) terms (adjusted *P* < 0.05) for time point- and organ-specific DEGs. Dot size reflects the number of genes per term, and color intensity indicates statistical significance. PI3K/PKB, phosphatidylinositol 3−kinase/protein kinase B; CX, cortex; MD, medulla; LV, liver; HR, heart; D7, day 7; D14, day 14; D21, day 21; and D35, day 35 time points. adj. p indicated adjusted *P* value. *n* = 6 male rats per group.

To further chareterized temporal tissue specific responses in SSH, we identified organ- and time-unique genes, defined as genes differentially expressed in only one organ at a single time point, regardless of their status at other stages. This set revealed dynamic and timed transcriptional changes across tissues (Figure 5C and Data sheet S6). To understand the biological impact of these dynamically regulated, organ- and time-specific gene sets, we performed GO-BP enrichment (Data sheet S7) and plotted the five most significantly (adjusted *P* < 0.05) enriched terms for each organ and time point (Figure 5D). While this focused view highlights the most prominent pathways at a specific time-point, additional significantly enriched terms were detected (Data sheet S7). This GO-BP analysis revealed that in the kidney cortex at day 7, the top pathways were involved in organic and monocarboxylic acid transport, pointing to an early metabolic-transport adjustment to high salt diet–induced hypertension. By day 21, top five GO-BP of the cortex shifted toward vascular and hemostatic regulation, with enrichment in pathways such as fibrinolysis, vasoconstriction, and proteolytic cascades. These features suggest the emergence of hemodynamic stress. The day 35 top five terms shift toward developmental and immune differentiation such as renal system development and lymphocyte differentiation, indicating late epithelial remodeling and immune engagement. Whereas in the kidney medulla, starting from day 7, enrichment was dominated by leukocyte activation and trafficking, indicating an early and robust inflammatory response. Terms related to catabolic and metabolic activities, such as amino acid and organic acid metabolism, rose to the top five pathways at later time points in the medulla. While the immune-related pathways did not appear among the top five terms at later stages, they remained significantly enriched in the medulla throughout the time course of study (Data sheet S7).

The liver displayed a shift from cell adhesion and focal adhesion at day 7 to lipid and fatty acid metabolic programs at days 21 and 35, including cellular ketone metabolism and fatty acid beta oxidation. These terms reflect the delayed but broad metabolic reprogramming seen in the liver during SSH. The heart showed little enrichment early on, with developmental terms on day 14. By day 35, prominent enrichment for contractile and ion transport programs emerged, including heart contraction, muscle system process, and metal ion transport, highlighting structural and functional remodeling in late-stage disease.

While our focused analysis of transient organ-time point responders revealed localized and stage-specific transcriptional programs, we next extended our analysis to all DEGs per organ and time point to capture broader patterns of biological pathways in salt-induced hypertension. We performed parallel GO-BP (Figure S7 and Data sheet S8) and Hallmark pathway enrichment analyses (Figure S8 and Data sheet S9) on the complete sets of DEGs, stratified by organ and time point. Together, these analyses reveal how salt-induced hypertension drives both temporal and organ-specific transcriptional responses, as well as broader biological processes that are active across multiple stages and shared among organs. By comparing these two levels of resolution, we show that individual organs deploy distinct molecular programs while also participating in common stress-related pathways. This integrated view helps explain how different tissues respond in parallel yet uniquely to SSH.

### Dynamic transcription factor-target gene networks coordinate specific responses in salt-induced hypertension

To infer upstream regulators of transcriptomic changes, we used the ChEA 2022 database, a high-confidence resource of transcription factor–target interactions from ChIP-seq and ChIP-chip studies^25,26^. Performing enrichment analysis of DEGs against the ChEA database identified candidate TFs likely driving the tissue- and time-specific transcriptional programs in SSH (Figure S9 and Data sheet S10). All TFs emerging from this analysis with adjusted *P* < 0.05 across all tissue and time point combinations were aggregated. From this comprehensive set, we further selected TFs that were themselves significantly differentially expressed, thereby prioritizing TFs both statistically enriched for targeting DEGs and directly responsive to SSH. This two-step filtering yielded a final set of 79 high-confidence TFs (Figure S10A). To investigate how TFs regulation evolves across tissues and over time, we applied both PCA and UMAP to the expression profiles of 79 high-confidence TFs (Figure S10B). PCA captured the major axes of transcriptional variance, with clear separation by tissue, reflecting distinct regulatory programs among analyzed tissues. In contrast, UMAP, which better preserves local structure, revealed tissue-specific expression patterns and gradual shifts across time points, offering a more detailed view of the evolving regulatory landscape during SSH progression (Figure S10B).

To identify which TFs differentiate tissue-specific responses over time, we applied random forest classifiers at each time point using the expression profiles of 79 TFs as input features and tissue identity as the classification target. Feature importance was assessed using the mean decrease in Gini impurity, a measure of how much each TF contributes to accurately distinguishing tissue-specific profiles. Through this supervised classification approach, we identified TFs most responsible for separating tissue-specific responses at different phases of SSH (Figure S10C). Next, to map the temporal involvement of TFs in response to SSH, we identify TFs that were significantly differentially expressed in one tissue at a given time point. These were classified as organ-time unique TFs capturing tissue and time restricted activity. A concentric donut plot (Figure 6A, Data sheet S11) visualized the distribution of these organ-time unique TFs across both tissue and time points. This analysis revealed tissue and time dependent heterogenity in TFs expression.

**Figure 6.**
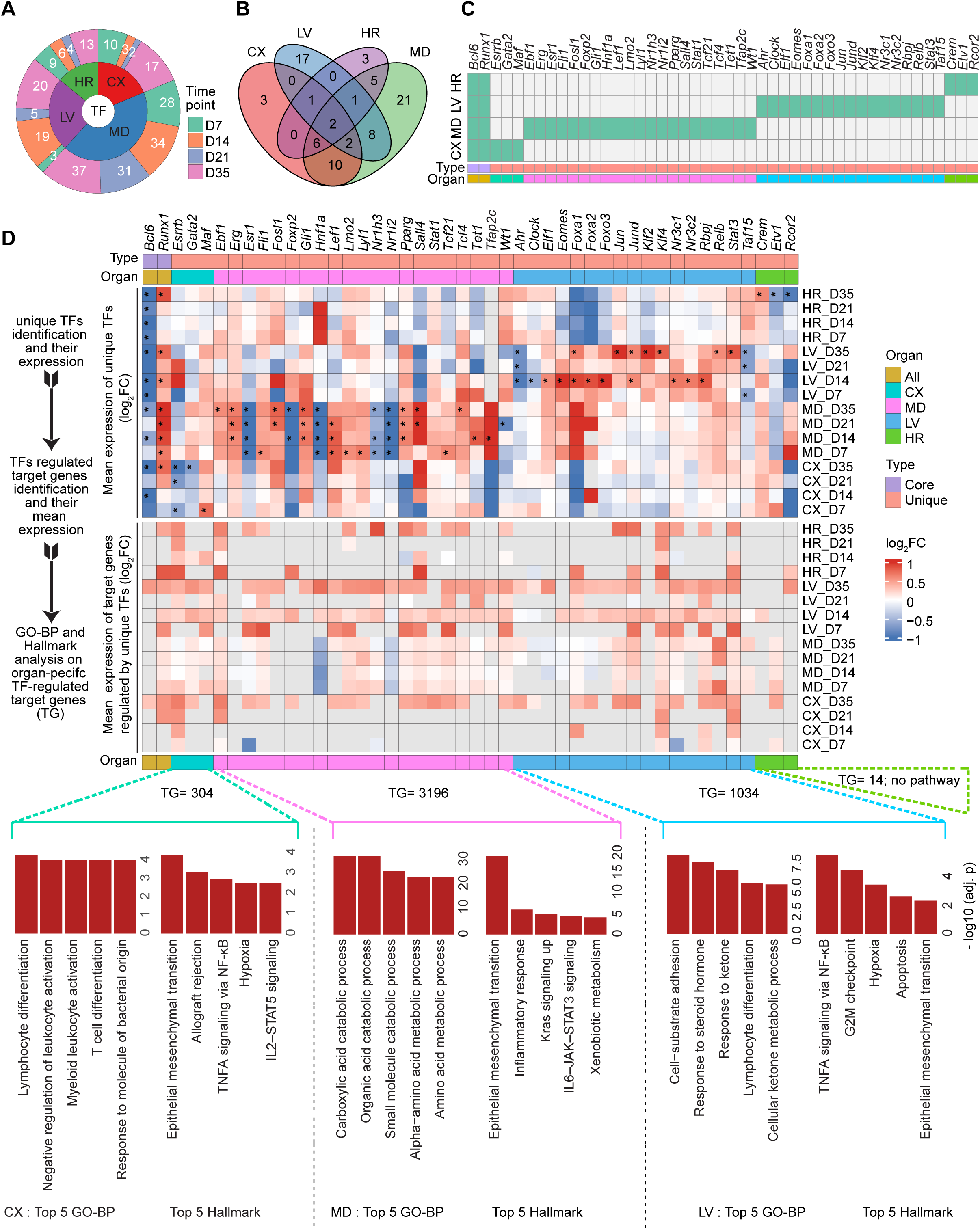
High salt diet-induced hypertension drives organ-specific regulatory programs. **A**. Concentric donut plot displaying the number of significantly differentially expressed transcription factors (TFs) per tissue (inner ring) and their distribution across four time points (days 7, 14, 21, and 35). **B**. Venn diagram depicting the overlap and exclusivity of TFs across cortex, medulla, liver, and heart, identifying shared versus tissue-specific TFs. **C**. Unique TFs per tissue and shared TFs visualized as a binary heatmap. **D**. TF-target genes network and functional relevance analysis. Top heatmap: heatmap showing the expression dynamics of two core and unique TFs across 16 tissue–time point combinations. Asterisks denote significantly altered TFs (|log₂ fold change| ≥ 0.585, equivalent to a fold change ≥ 1.5 and adjusted *P* < 0.05); Middle heatmap: showing mean log₂FC of TF-regulated target genes; Lower pathway panel: showing top 5 GO Biological Processes (left) and Hallmark pathways (right) for CX, MD, and LV emerged by TF regulated target genes. Bar heights represent –log₁₀ adjusted *P* values. Target gene (TG) counts indicate the number of organs-specific genes regulated by corresponding TFs and used in the pathway analysis. No pathways met significance thresholds in HR. Enriched pathways with adjusted *P* < 0.05 were considered significant. CX, cortex; MD, medulla; LV, liver; HR, heart; D7, day 7; D14, day 14; D21, day 21; and D35, day 35 time points. adj. p indicated adjusted *P* value. *n* = 6 male rats per group.

To assess the degree of overlap in transcriptional regulation across tissues, we constructed a four-way Venn diagram of all TFs (Figure 6B). This analysis revealed 44 TFs exhibiting organ-specific regulation. Only two TFs, *Bcl6* and *Runx1*, were shared across all four tissues, indicating that TF activity in SSH is largely tissue-specific. To explore this organ specificity in greater detail and extend the overlap patterns seen in the Venn diagrams, we generated a binary presence–absence heatmap of organ-unique TFs (Figure 6C), providing a more granular view by displaying each TF individually, and revealing clear segregation of TFs by tissues. Next, to assess whether organ-unique TFs impose functional control on downstream genes, we mapped ChEA predicted targets for each of the 44 TFs that remained restricted to a single tissue across the experiment. For every tissue, we compiled only the targets linked to its own unique TFs and overlaid TFs and target gene expression onto our RNA-seq data. The expression dynamics of multiple organ-unique TFs paralleled those of their predicted targets, indicating the functional coherence of these regulatory relationships (Figure 6D). This coordinated activity suggests that TFs do not operate in isolation but instead regulate structured transcriptional modules that drive specific responses in SSH. The magnitude of these regulons varied substantially across tissues, with medulla-unique TFs regulating the largest gene networks, followed by those in the liver, cortex, and heart. pathway analysis of the organ-specific target pools reveals distinct biological themes. Cortex regulons were enriched for immune cell differentiation and activation, EMT, TNFA signaling via NF-κB, hypoxia, and IL2–STAT5 signaling. Alongside, medulla regulons were dominated by catabolic and metabolic processes, Kras signaling, IL6–JAK–STAT3 signaling and xenobiotic metabolism. Whereas, liver-restricted TFs were associated with pathways involving cell-substrate adhesion, steroid hormone response, ketone metabolism, lymphocyte differentiation, TNFA signaling via NF-κB, hypoxia, apoptosis, and EMT (Figure 6D and Data sheet S12).

Together, we identified 79 high-confidence TFs and mapped their dynamic, tissue-specific regulatory networks in SSH, yielding a curated list of candidate targets for mechanistic and translational studies.

### Integration with the human genome-wide association studies (GWAS) reveals a conserved hypertension- and CKD-linked transcriptional response

To determine whether the transcriptional programs altered in our hypertensive rat model are relevant to human disease, we integrated our transcriptomic data with human GWAS loci for the terms “hypertension” and “chronic kidney disease”. In this analysis, first DEGs from the kidney cortex, medulla, liver, and heart were mapped to their human orthologs. Among the 5,213 rat DEGs mapped to human orthologs, 100 genes overlapped with hypertension-associated GWAS genes, comprising a total of 386 genes (Figure 7A), and 143 genes intersected with CKD-associated genes spanning 406 total genes (Figure 7B). These overlaps represented highly significant enrichments (Fisher’s exact test, p < 0.0001; odds ratios = 2.4 for hypertension and 3.8 for CKD), highlighting that the transcriptional shifts seen in our rat model overlap strongly with known human hypertension and CKD risk loci-associated genes. Expression analysis of these overlapping genes revealed distinct temporal and tissue-specific expression patterns (Figure S11). Further analysis across GO-BP, KEGG, and Reactome databases shows that hypertension-linked genes are dominated by developmental and endocrine pathways. Top terms include epithelial morphogenesis, renal system development, hormone metabolism, renin and cortisol secretion and extracellular-matrix organization (Figure 7C, Figures S12A–C). In contrast, CKD-linked genes were enriched for metabolic and detoxification pathways (Figures 7D and S12A, D, and E).

**Figure 7.**
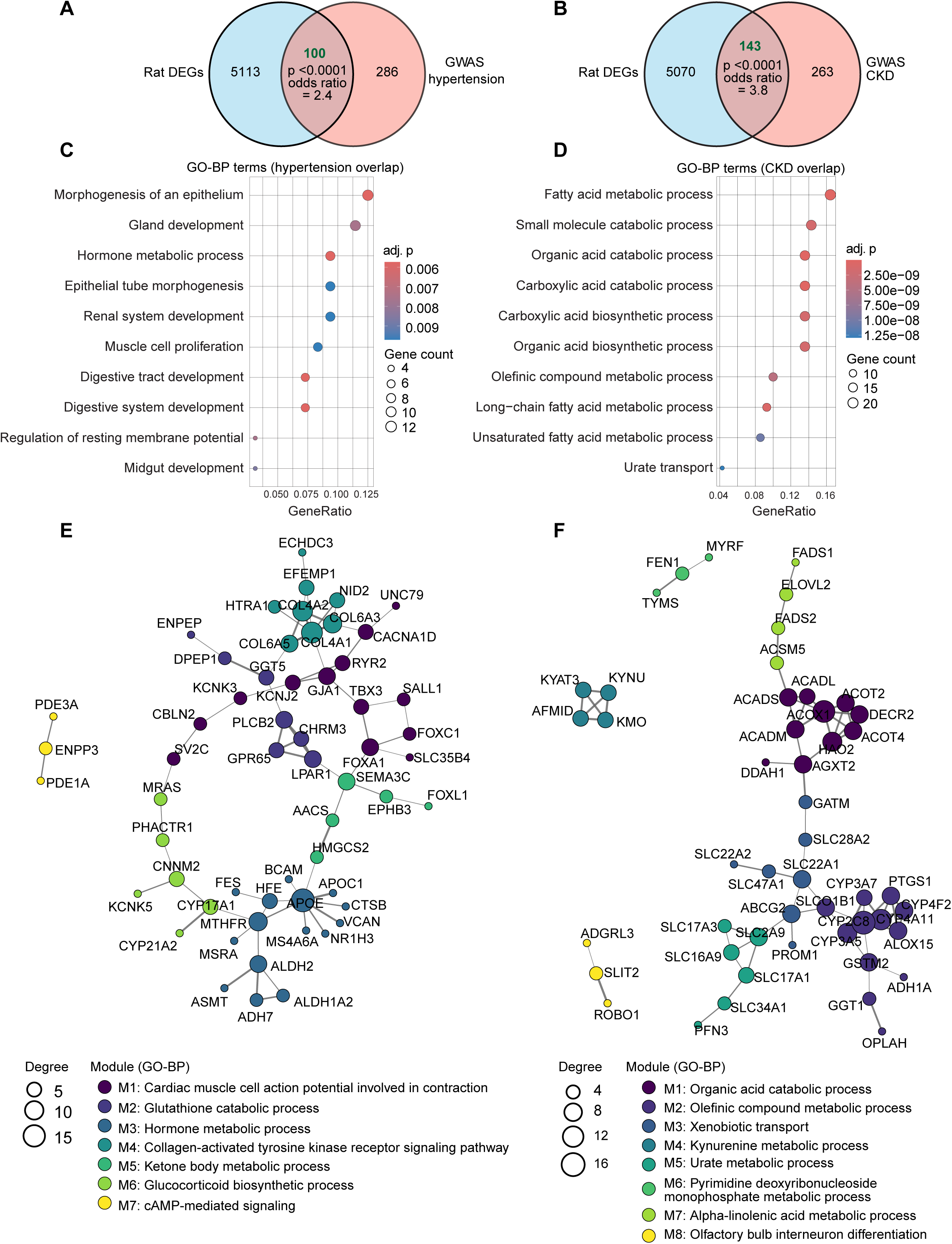
Integration of Dahl SS rat transcriptomes with human genome-wide association studies (GWAS) loci identifies conserved gene networks underlying hypertension and kidney disease. **A** and **B**. Integration of rat transcriptomic data with human GWAS identified a significant enrichment of overlap genes associated with hypertension (**A**) and chronic kidney disease (CKD; **B**). Venn diagrams show the intersection between rat DEGs and GWAS loci associated genes, revealing non-random convergence (hypertension: *P* < 0.0001, odds ratio = 2.4; CKD: *P* < 0.0001, odds ratio = 3.8; hypergeometric test). **C** and **D**. Enriched pathways from GWAS-overlapping genes revealed coherent biological processes in hypertension (**C**), and CKD-overlap genes (**D**). **E** and **F**. Protein-protein interaction (PPI) networks, constructed from GWAS-overlap genes, revealing distinct modular architectures (STRING: score > 400 for hypertension, > 700 for CKD). Louvain clustering identified discrete gene communities, each annotated by its most significantly enriched GO-BP term. Node size corresponds to degree centrality; edge width reflects interaction strength; colors denote module identity. adj. p indicated adjusted *P* value.

To translate the GWAS-overlap lists into mechanistic circuitry, we constructed PPI networks using STRING (confidence > 0.4 for hypertension and > 0.7 for CKD) and applied Louvain clustering^27^ to identify discrete communities. After mapping, singletons were removed, and the remaining interaction graphs were partitioned with the Louvain algorithm. Communities containing two or fewer proteins were excluded in order to retain robust modules. To infer the biological impact of community modules, we ran GO-BP enrichment on communities and labelled modules with their top biological process term. With this approach, 100 GWAS overlap gene-associated proteins of hypertension were segregated into seven modules (Figure 7E). A connected ion-flux core (M1, cardiac muscle cell action potential involved in contraction) was anchored by calcium-handling hubs *calcium voltage-gated channel subunit alpha1 D* (*CACNA1D*) and *potassium inwardly rectifying channel subfamily J member 2* (*KCNJ2*), together with the gap-junction protein *gap junction protein alpha 1* (*GJA1*), defining an electrically responsive scaffold. A collagen-receptor module (M4), built around *collagen type IV alpha chain* (*COL4A*) and *collagen type VI alpha 3 chain* (*COL6A3*), linked extracellular matrix remodeling to this ion-flux hub, consistent with pressure-induced structural adaptation. The hormone-metabolism module (M3) expanded this circuitry by incorporating one-carbon and lipid transport elements, such as *Methylenetetrahydrofolate reductase* (*MTHFR*), which aligns folate-dependent methyl flux with the cholesterol carrier *Apolipoprotein E(APOE*) and a suite of steroid-modifying enzymes. *MTHFR* also maintained a high-confidence connection to c*ytochrome P450 family 17 subfamily A member 1* (*CYP17A1*), the anchor of the steroid-biosynthetic module (M6). This bridge places methyl-group supply in close proximity to cortisol and androgen synthesis, suggesting that folate status could influence endocrine output during the hypertensive response. In contrast, the cAMP-signaling triad (M7), containing *phosphodiesterase 3A* (*PDE3A*), *phosphodiesterase 1A* (*PDE1A*), and *ectonucleotide pyrophosphatase/phosphodiesterase 3 (ENPP3*), was topologically isolated, indicating that cyclic-nucleotide regulation operates as an independent signaling node.

Analysis of the CKD GWAS-overlap interactome (Figure 7F) resolved eight discrete communities, illustrating how distinct metabolic programs are organized and coordinated. The M1 module, annotated as organic-acid catabolic process, represents a β-oxidation axis anchored by *acyl-CoA dehydrogenase long chain* (*ACADL*), *acyl-CoA dehydrogenase medium chain* (*ACADM*), *acyl-CoA dehydrogenase short chain* (*ACADS*), and the peroxisomal oxidase *acyl-CoA oxidase 1*(*ACOX1*). Attached to M1, the M2 module (olefinic-compound metabolic process) includes the long-chain desaturases *fatty acid desaturase 1* (*FADS1*), *fatty acid desaturase 2* (*FADS2*), and the elongase *ELOVL fatty acid elongase 2* (*ELOVL2*), showing that unsaturated fatty acid synthesis feeds into downstream β-oxidation. A multifunctional detoxification cluster emerged alongside these lipid modules, enriched for cytochrome P450 enzymes, the eicosanoid synthase *arachidonate 15-lipoxygenase* (*ALOX15*), and glutathione-handling proteins *glutathione S-transferase mu 2* (*GSTM2*) and *gamma-glutamyltransferase 1* (*GGT1*). Edges connecting adjacent transporter-enriched communities, comprising SLC family members *solute carrier family 22 member 1* (*SLC22A1*)*, solute carrier family 22 member 2* (*SLC22A2*)*, solute carrier family 2 member 9* (*SLC2A9*)*, solute carrier family 28 member 2* (*SLC28A2*) and the ABC transporter *ATP binding cassette subfamily G Member 2* (*ABCG2*), suggest potential coordination in handling lipid-derived metabolites and purine catabolites through parallel excretory pathways. In parallel, several metabolic programs remained topologically insulated. A compact kynurenine module composed of *kynureninase* (*KYNU*), *kynurenine 3-monooxygenase* (*KMO*), *kynurenine aminotransferase 3* (*KYAT3*), and *arylformamidase* (*AFMID*) forms a dedicated tryptophan-metabolism circuit. Likewise, the nucleotide metabolism group, including (*Flap structure-specific endonuclease 1* (*FEN1*), *thymidylate synthase* (*TYMS*), and *myelin regulatory factor* (*MYRF*), and the developmental cluster, comprising *roundabout guidance receptor 1* (*ROBO1*), s*lit guidance ligand 2* (*SLIT2*), and *adhesion G protein-coupled receptor L3* (*ADGRL3*), occupy peripheral positions.

In summary, our data recapitulate gene expression patterns linked to human GWAS loci for hypertension and CKD, including key disease pathways reported in humans. These conserved signatures underscore the translational relevance of our findings for dissecting multi-organ disease mechanisms in hypertension.

### Compound–transcriptome mapping reveals stage- and organ-specific therapeutic opportunities

One of the most powerful applications of transcriptomic predictive analysis lies in identifying pharmacological targets through gene remodeling patterns. Leveraging our longitudinal data, we aim to map these transcriptional changes onto pharmacological signatures to guide therapeutic strategies for controlling and reversing hypertension. To systematically identify compounds that could reverse SSH-associated transcriptional programs, we leveraged the Library of Integrated Network-Based Cellular Signatures (LINCS) database. Developed through the National Institutes of Health (NIH) Common Fund LINCS program, this resource provides a comprehensive catalog of gene expression and cellular responses to thousands of small molecules and genetic perturbations^28,29^.

Our LINCS-based predictive compound–transcriptome analysis identified several high-confidence compounds with the potential to reverse maladaptive gene profiles across organs. In the kidney cortex, NVP-BEZ235 (dual phosphoinositide 3-kinase/mechanistic target of rapamycin [PI3K/mTOR] inhibitor), dovitinib (fibroblast growth factor receptor/vascular endothelial growth factor receptor [FGFR/VEGFR] inhibitor), and palbociclib (cyclin-dependent kinase 4/6 [CDK4/6] inhibitor) were predicted to most effectively counter sustained induction of E2F target genes, G2M checkpoint components, and mechanistic target of rapamycin complex 1 (mTORC1) signaling pathways driving proliferative expansion and metabolic alterations (Figure 8A). These findings suggest that coordinated inhibition of growth factor receptors and cell-cycle regulators may restore transcriptional homeostasis in cortical tissue.

**Figure 8.**
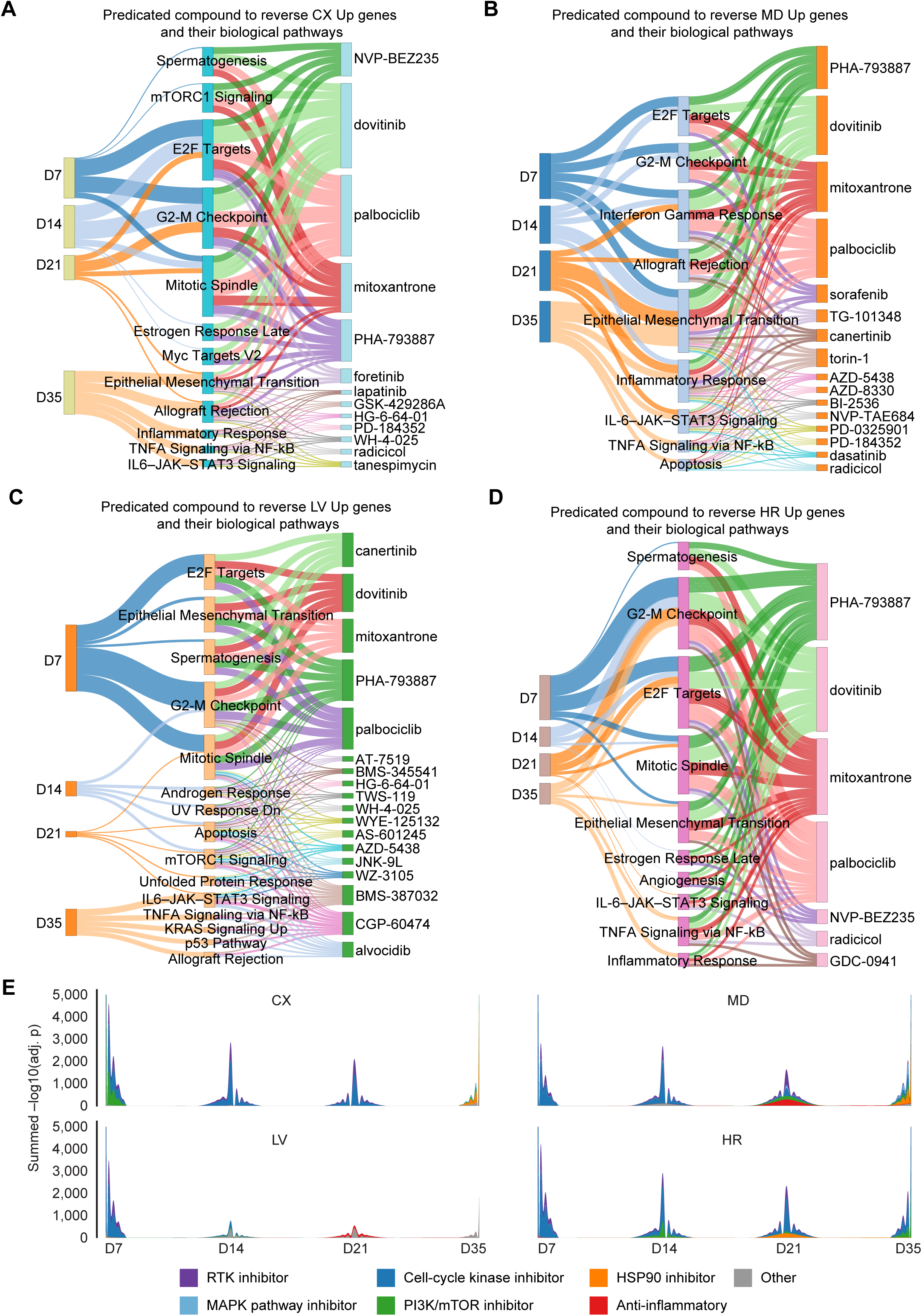
Temporal and organ-specific prediction of small-molecule modulators to target salt-sensitive hypertension. **A–D**. Sankey diagrams integrating differential expression profiles with LINCS L1000 chemical perturbation signatures for significantly upregulated genes in the kidney cortex (**A**), kidney medulla (**B**), liver (**C**), and heart (**D**). Each network connects time points (days 7–D35; left), enriched top 5 Hallmark pathways (middle), and the top predicted small molecules (right) prioritized for their potential to reverse disease-associated transcriptional changes. Edge thickness denotes −log₁₀(adjusted *P* value), indicating enrichment strength. **E**. Stacked stream graphs show, for each tissue, the cumulative enrichment of small-molecule classes at each time point, represented as the summed – log₁₀(adjusted *P* value) of all compounds within each class. Peaks indicate phases when a given class is most strongly aligned with the organ-specific transcriptome. CX, cortex; MD, medulla; LV, liver; HR, heart; D7, day 7; D14, day 14; D21, day 21; and D35, day 35 time points. adj. p indicated adjusted *P* value. *n* = 6 male rats per group.

In the kidney medulla, PHA-793887 (CDK inhibitor), dovitinib, mitoxantrone, and sorafenib (multi-kinase inhibitor) emerged as prominent candidates predicted to reverse maladaptive proliferative programs and concurrent activation of interferon-gamma and Janus kinase/signal transducer and activator of transcription 3 (JAK/STAT3) inflammatory pathways (Figure 8B). Notably, selective mitogen-activated protein kinase kinase (MEK) inhibitors, including PD-0325901 and PD-184352, were also associated with reversal of downregulated metabolic pathways such as fatty acid oxidation and oxidative phosphorylation (Figures S13A and B), suggesting their potential to reactivate suppressed metabolism. This combinatorial signature highlights a dual proliferative-inflammatory axis as therapeutic target in medullary injury.

In the liver, compounds such as canertinib (epidermal growth factor receptor/human epidermal growth factor receptor 2 [EGFR/HER2] inhibitor), torin-1 (mTOR inhibitor), palbociclib, and CGP-60474 (broad-spectrum CDK inhibitor) were consistently prioritized to counter maladaptive proliferation, unfolded protein response, and NF-κB-mediated inflammatory signaling (Figure 8C). Additionally, MEK inhibitors (PD-0325901, AZD-8330, and selumetinib), along with decitabine (a DNA methyltransferase inhibitor) and withaferin A (a natural steroidal lactone), were predicted to restore downregulated transcriptional programs involving fatty acid metabolism, mTORC1 signaling, peroxisome activity, and cholesterol homeostasis (Figure S13C). These findings underscore the dual potential of kinase and epigenetic modulators to suppress pathogenic activation while reactivating metabolic pathways central to hepatic remodeling.

In the heart, enrichment was observed for PHA-793887, dovitinib, mitoxantrone, NVP-BEZ235, radicicol (heat shock protein 90 [HSP90] inhibitor), and GDC-0941 (PI3K inhibitor) to counter early mitotic gene induction and subsequent angiogenic and inflammatory activation (Figure 8D). MEK inhibitors (trametinib, AZD-8330), EGFR/HER2 inhibitors (neratinib, erlotinib, afatinib), and radicicol were recurrently associated with restoration of downregulated programs spanning xenobiotic metabolism, adipogenesis, and estrogen response (Figure S13D), highlighting the capacity of pathway-selective inhibitors to normalize both hyperactivated and suppressed networks.

To further resolve how pharmacologic predictions change over the course of disease, we classified compounds into mechanistic classes and quantified their enrichment dynamics across disease progression (Figure 8E). Temporal analysis revealed early enrichment of cell-cycle and receptor tyrosine kinase (RTK) inhibitors, shifting toward PI3K/mTOR and HSP90 inhibitors at later stages. These patterns demonstrate that different classes of compounds are preferentially predicted at distinct disease phases, further reflecting evolving pathophysiological mechanisms as SSH progresses. Collectively, this integrated analysis delineates pharmacologic candidates and their impact on different biological pathways. These findings highlight the need for precision medicine approaches. Such approaches should not focus solely on lowering blood pressure but also account for the duration of hypertension, organ damage, and associated molecular pathways.

## DISCUSSION

In this study, we demonstrate that salt-sensitive hypertension progresses through a conserved early proliferative program, followed by divergent immune and fibrotic remodeling across organs. The kidney medulla exhibits the most pronounced and sustained changes, marked by robust inflammatory activation and metabolic suppression, whereas the cortex, liver, and heart show delayed, tissue-specific programs. We identified 79 high-confidence regulators, largely with organ-restricted activity, with *Bcl6* and *Runx1* emerging as the major conserved regulators across tissues. Integration with human GWAS loci underscores the translational relevance of our findings, linking experimental SSH signatures to established hypertension- and CKD-associated genes and pathways. Compound–transcriptome mapping identified stage- and organ-specific therapeutic opportunities. Inhibitors of PI3K/mTOR, CDKs, and RTKs emerged as candidates for early disease stages, whereas later stages could be better ameliorated by modulators of MAPK signaling, HSP90 inhibitors, and agents targeting epigenetic regulation.

The renal medulla has long been recognized as particularly vulnerable to injury because of low oxygen gradients, significant sodium transport activity, and relative hypoperfusion^10,30^. Hypoxia and oxidative stress have been implicated as key drivers of injury in this region, yet the molecular pathways linking these physiological stressors to progressive damage remain poorly defined. Our longitudinal transcriptomic analysis helps close this gap. Our data indicate that medullary susceptibility in SSH is manifested by the early and sustained activation of immune and proliferative programs, with metabolic suppression, thereby establishing conditions that promote progressive kidney injury. Notably, because albuminuria emerges as one of the earliest detectable manifestations in SSH, cortical injury is often considered central to disease progression. In earlier work, we demonstrated that cortical glomeruli and tubules undergo metabolic dysregulation, oxidative stress, and structural remodeling during SSH^20^. Nevertheless, it remained unresolved whether cortical alterations or the medullary injury are the major events in the SSH-driven CKD. Here, we provide a direct comparison between the cortex and medulla within the kidney. We found that, despite their proximity, the cortex showed a delayed and transient transcriptional programs, with a brief rise in metabolic activity followed by later immune activation. This pattern contrasted with the medulla, which showed early cast formation, cellular injury, and fibrosis, while cortical injury remained comparatively moderate. Functionally, albuminuria rose early and later slightly declined, reflecting an initial phase of hyperpermeability or hyperfiltration followed by nephron loss. Serum creatinine also increased only at a late stage, aligning with the cortical damage observed in later stages in histological analysis. Together, our longitudinal transcriptomic, histological, and functional data provide important clarity that the cortex initially shows a metabolically adaptive phenotype characterized by increased oxidative metabolism and glomerular hyperfiltration. Over time, however, this adaptive state shifts toward immune activation and extracellular matrix remodeling. Whereas, the medulla emerged as the primary site of injury and possibly a major driver of CKD in SSH. Our study also underscores the limited sensitivity of conventional serum creatinine measurements for detecting early SSH-induced kidney injury. This highlights the need for more comprehensive biomarkers capable of capturing both alterations in filtration and early medullary stress before irreversible damage occurs.

Beyond the kidney, emerging evidence suggests that SSH also engages the liver, positioning it as an important yet underappreciated contributor to disease progression. The liver is increasingly recognized as an active modulator of systemic hypertension, metabolic dysfunction, and CVD^31,32^. Notably, epidemiological data indicate that high salt intake is linked to a greater risk of developing nonalcoholic fatty liver disease and advanced liver fibrosis^33–36^. Furthermore, experimental studies on murine models indicate that a high salt diet induces epigenetic modifications in the liver that sustain hepatic steatosis and inflammation, ultimately contributing to cardiovascular injury^13^. Within this evolving concept, our analysis provides a longitudinal view of hepatic transcriptional remodeling in SSH. Our findings indicate that the liver actively responds to salt-induced hypertension thorugh metabolic and inflammatory signals, which may contribute to vascular dysfunction, disrupted lipid homeostasis, and systemic inflammation. Our results emphasize that SSH is not solely a renal or cardiovascular disease but also involves dynamic hepatic changes that may amplify cardiovascular risk through distinct molecular pathways and needs more detailed investigations.

The cardiac consequences of SSH have historically been attributed to chronic pressure overload and neurohumoral activation^37–39^. However, accumulating evidence suggests that local metabolic and inflammatory remodeling contribute independently to cardiac hypertrophy and dysfunction^40–43^. Our study reveal that the heart mounts an early proliferative stress response, which transitions into inflammatory and fibrotic remodeling, pointing to intrinsic molecular triggers rather than just mechanical load. This sequence mirrors earlier observations of hypertrophic cardiomyocyte signaling gradually shifting into maladaptive inflammation and tissue remodeling^44–46^. Notably, our findings highlight IL6–JAK–STAT3 as a central non-hemodynamic mediator in the heart’s response to SSH^47^. This is consistent with preclinical models where IL-6 deficiency impairs angiotensin II–induced hypertension and cardiac remodeling^48^. Activation of JAK–STAT3 in this context is known to amplify cardiac inflammation and fibrosis even in the absence of sustained pressure overload^49^. Together, these insights suggest that SSH engages both mechanical and immune–metabolic pathways in the heart, with STAT3-dependent signaling emerging as a potential driver of maladaptive remodeling. Targeting this axis at the critical transition from proliferative stress to inflammatory remodeling may represent a promising strategy to prevent progression to heart failure in SSH.

Along with organ-specific responses, the identification of the shared proliferative program across organs is noteworthy, as most prior work has emphasized immune and fibrotic remodeling as central drivers of hypertensive kidney damage. Our data suggest that SSH initially elicits a coordinated proliferative response, which may reflect an adaptive attempt at tissue repair. Similar early activation of cell cycle networks has been observed in kidney injury^50,51^ and cardiac hypertrophy^52^. However, when this response is inadequately controlled or sustained by long-term stress, it often precedes the transition to maladaptive remodeling^53,54^. We found that in SSH, a shared proliferative phase shifts to organ-specific immune and extracellular-matrix programs. Our longitudinal, multi-organ atlas traces this divergence toward organ-specific pathology and highlights its importance for more studies.

At the level of transcriptional regulation, beyond the identification of tissue-specific regulators, we identified *Bcl6* and *Runx1* as opposing master regulators, revealing a unifying mechanistic axis that integrates proliferative, inflammatory, and fibrotic responses across organs. *Bcl6* has been shown as suppressor of NF-κB-dependent cytokine expression, with studies showing that its overexpression ameliorates renal, hepatic, and vascular inflammation^55–58^. Conversely, *Runx1* has emerged as a critical driver of proinflammatory and profibrotic gene expression across multiple cell types, including macrophages, vascular smooth muscle cells, and tubular epithelial cells^59,60^. Our observation of consistent *Bcl6* downregulation and *Runx1* upregulation across organs suggests a coordinated transcriptional switch that may be exploited therapeutically in SSH. Of note, *Runx1* inhibition has been shown to be beneficial for pulmonary arterial hypertension and myocardial infarction^61–63^, although its role in SSH and associated multi-organ damage remains unexplored.

Integrating transcriptomic signatures with human GWAS data allowed us to examin our findings through the lens of established genetic risk loci for hypertension and CKD. This approach not only strengthens the biological relevance of our experimental findings but also points to evolutionarily conserved pathways that may shape disease susceptibility. In this context, folate metabolism (*MTHFR*), steroidogenesis (*CYP17A1*), and extracellular matrix regulation (*COL4A1, COL6A3*) aligned with hypertension risk, while fatty acid oxidation (*ACADL, ACOX1*), solute transport (*SLC22A1/2*), and tryptophan metabolism (*KYNU, KMO*) were linked to CKD risk. Among these, *MTHFR* is particularly notable, pointing to conserved one-carbon metabolism, which is not fully understood in SSH. *MTHFR* encodes methylenetetrahydrofolate reductase, a critical enzyme that regulates homocysteine–methionine balance and supplies methyl groups for DNA and histone methylation, thereby shaping epigenetic patterns that regulate vascular tone, renal sodium handling, and inflammatory signaling^64^. Dysregulation of *MTHFR* has been associated with endothelial dysfunction and increased susceptibility to CVD^65,66^, yet their relevance to SSH has not been systematically explored. Similarly, the kynurenine arm of tryptophan metabolism contributes to immunoregulation, redox balance, and NAD⁺ biosynthesis^67^. Previously, we have shown the protective role of the kynurenine pathway against ischemic acute kidney injury in mice^68^. However, in the Dahl SS rat model, and especially in the settings of SSH and associated CKD, the role of the kynurenine pathway is not well characterized. These conserved mechanisms underscore the translational significance of our findings and position metabolic–immune cross-talk as a central driver of SSH progression.

Organ damage in SSH often progresses despite intensive blood pressure control. High sodium itself promotes vascular dysfunction, oxidative stress, and fibrosis, including arterial stiffening via profibrotic mediators such as transforming growth factor-beta (TGF-β), even in normotensive conditions^69–71^. These findings indicate that SSH-induced injury is not simply a secondary consequence of elevated blood pressure but reflects parallel molecular processes that require targeted therapeutic strategies. Preclinical studies have demonstrated that interventions directed at hypertrophy, inflammation, or fibrosis can mitigate organ damage, yet these benefits are often highly context dependent and diminish when therapies are applied outside the optimal disease window^72–77^. Our pharmacotranscriptomic analysis provides an explanation for this variability by showing that SSH activates temporally distinct and organ-specific transcriptional programs. Early proliferative responses transition into immune activation and fibrotic remodeling, shifting therapeutic opportunities across time and tissue compartments. These dynamics suggest that drugs effective in one phase may lose efficacy if applied at another. Clinically, this suggests that therapy for SSH cannot be determined by blood pressure control alone but must also take into account the evolving molecular and functional state of the organs. Improved biomarkers that capture organ-specific trajectories are therefore needed to guide intervention timing and choice. Together, these insights argue for stage- and organ-tailored therapeutic strategies that move beyond conventional one-size-fits-all blood pressure management.

Despite its strengths, this study has limitations. We used mRNA sequencing to achieve high coverage and statistical power to generate a detailed transcriptomic view of SSH. However, this approach cannot resolve cell type–specific transcriptional heterogeneity. We prioritized depth and sensitivity, which remain limited with current single-cell technologies. Recently, recognizing the importance of cellular resolution, research efforts have initiated single-cell mapping of hypertension^78^. More studies using single-cell and spatial transcriptomics will be essential to define both cell type- and time-specific contributions with greater precision. Additionally, although the Dahl SS rat is a well-established model of human SSH, which is also supported by our GWAS analysis, species differences should be considered when translating these findings to humans.

In summary, our study presents a transcriptomic atlas of SSH progression across kidney, liver, and heart, which can be leveraged to explore diverse mechanistic pathways.

## Acknowledgments

The authors thank the staff of the Division of Comparative Medicine (DCM) at the University of South Florida for their dedicated care and support of the animals used in this study. Their expertise in managing the animal housing facility and ensuring high standards of animal welfare was essential for the successful completion of this research.

## Author contributions

R.T. and A.S. conceptualized and designed the study. R.T., V.L., and M.L. performed the experiments and collected samples. R.T., O.K., L.V.D., M.L., B.X., R.B., S.D., D.V.I., and O.P. analyzed the data. R.T. and A.S. interpreted the data. R.T. wrote the manuscript and finalized it for publication. All authors reviewed and approved the final version of the article.

