## Supplemental Materials for "Longitudinal Multi-Organ Transcriptomic Atlas of Salt-Induced Hypertension"

**Running title:** Transcriptomic Atlas of Salt-Induced Hypertension

#### MATERIAL AND METHODS

##### Animals

Male Dahl salt-sensitive (SS/JrHsdMcwi) rats were used as a model of salt-induced hypertension. Animals were housed in the AAALAC-accredited animal facility at the University of South Florida under specific pathogen-free conditions, maintained on a 12-hour light/dark cycle in temperature- and humidity-controlled rooms, with *ad libitum* access to food and water.

Dahl SS rats were maintained on a normal-salt diet control (NS, 0.4% NaCl, # D113755, Dyets Inc). To induce salt-sensitive hypertension, 9–10-week-old animals were switched to a high-salt diet (HS, 4% NaCl , # D113756, Dyets Inc) for 7, 14, 21, or 35 days. Animals on normal-salt diet were used as control group. One day before the final day of selected time points (day 7, 14, 21, and 35), rats were individually housed in metabolic cages for 24-hour urine collection. At each time-point, animals were euthanized under deep isoflurane/$O_2$ anesthesia and prior to renal perfusion, heparinized blood was obtained from the descending aorta and centrifuged at 5,000 g for 5 minutes to isolate plasma. Kidney cortex, kidney medulla, liver, and heart tissues were snap-frozen in liquid nitrogen or fixed in formalin for transcriptomic and histological assessments. Blood and urine samples were used for biochemical profiling. All experimental procedures were approved by the Institutional Animal Care and Use Committee (IACUC) at the University of South Florida.

##### Blood pressure measurement

We monitored blood pressure for 21 days on a different cohort of Dahl rats using the DSI telemetry system to confirm the blood pressure elevation with HS diet and establish the hypertensive phenotype. Male Dahl SS rats were anesthetized with 2–3% (vol/vol) isoflurane, and a blood pressure transmitter (PA-C40; DSI) was implanted subcutaneously, with the catheter tip positioned in the abdominal aorta via the femoral artery. After a 3-day recovery, blood pressure in conscious, unrestrained rats was continuously recorded under NS and HS diet conditions as described previously[1,2].

##### RNA Sequencing

Total RNA was extracted from snap-frozen kidney cortex, kidney medulla, liver, and heart tissues using TRIzol™ Reagent (Invitrogen). Sample were submitted to Novogene Corporation Inc for RNA sequencing. Briefly, RNA quality was assessed using a Bioanalyzer 2100 (Agilent Technologies). Only high-quality RNA (RNA Integrity Number (RIN) = 8.7 ±0.9) was used for library preparation. mRNA was purified using poly-T oligo-attached magnetic beads and subsequently fragmented. First-strand cDNA synthesis was performed using random hexamer primers, followed by second-strand synthesis. The library was prepared and checked with Qubit and real-time PCR for quantification and bioanalyzer for size distribution detection. After quality check, libraries were pooled and sequenced on an Illumina platform using paired-end sequencing using the Illumina "Sequencing by Synthesis" (SBS) chemistry. Raw data of fastaq format was cleaned and used for downstream analysis. Clean, paired-end reads were aligned to the *Rattus norvegicus* reference genome (Ensembl Rnor_6.0) using HISAT2 v2.2.1. Gene-level counts were computed using *featureCounts* (v2.0.6) and normalized as FPKM (Fragments Per Kilobase of transcript per Million mapped reads) for expression profiling. Differential gene expressions between conditions were analyzed using DESeq2 for datasets with biological replicates. Genes with an $|\log_2$ fold change$| \geq 0.585$, equivalent to a fold change $\geq 1.5$, adjusted $P < 0.05$ were considered significantly differentially expressed genes (DEGs).

#### Transcriptomic analysis

Analyses were conducted in R studio (v4.3.1) using DESeq2, tidyverse, ComplexHeatmap, msigdbr, enrichR, STRINGdb, and other required packages. Counts were processed using the variance-stabilizing transformation (VST) in DESeq2 for unsupervised dimensionality reduction and visualization. Principal component analysis (PCA) was applied using the prcomp function, and the first two components were plotted to assess organ- and timepoint-specific separation. Sample–sample Pearson correlations were computed on the full VST matrix and visualized as annotated heatmaps using pheatmap. To capture non-linear structure, uniform manifold approximation and projection (UMAP) embeddings were generated (uwot, n_neighbors = 15, min_dist = 0.1) and visualized with ggplot2. Volcano plots were generated using counts per gene ID per organ–timepoint combination, with upregulated, downregulated, and nonsignificant genes distinguished by color. For each comparison, DEGs were intersected across tissues and timepoints to identify shared and unique responses. Common DEGs were visualized using Venn diagrams based on gene symbols (VennDiagram), clustered heatmaps (pheatmap), and subjected to Gene Ontology Biological Process (GO–BP), Kyoto Encyclopedia of Genes and Genomes (KEGG), and Hallmark enrichment via clusterProfiler and msigdbr.

#### Pathway activity scoring and trajectory analysis

Transcriptome-wide pathway activity was quantified using 50 Hallmark gene sets (MSigDB v7.5.1). Pathway scores were computed as the mean $\log_2$FC for all genes in each set across organ–timepoint combinations, yielding a 50×16 matrix. These scores were visualized as heatmaps, ridge plots (ggridges), and Euclidean distance trajectories relative to baseline to capture global shifts. Inter-organ pathway similarity was evaluated using pairwise Pearson correlation of pathway vectors and displayed as correlation heatmaps. Hierarchical clustering (Ward's method, Euclidean distance) identified stable modules of co-regulated pathways, with cluster identity and biological function annotated. UMAP and multidimensional scaling (MDS) were applied to visualize higher-order pathway dynamics. Organ-specific patterns were highlighted by extracting the top 10 most variable pathways per tissue and plotting temporal trajectories. Directionality of pathway regulation (up vs. down) was summarized using stacked bar plots across time.

#### Protein-protein interaction analysis

To interrogate gene-level convergence, protein-protein interaction (PPI) networks were constructed for common DEGs at early (D7) and late (D35) timepoints using the STRING database[3] (v11, score > 700). Networks were clustered using Louvain community detection[4] (igraph) and annotated via module-level enrichment against Hallmark gene sets. Visualizations were rendered with ggraph, where nodes represent genes sized by connectivity and colored by module identity.

#### Upstream transcription factor and network analysis

Transcription factor (TF) enrichment was performed using ChEA 2022[5] via the enrichr R interface. To enable compatibility with human TF databases, rat DEGs were mapped to human orthologs using a tiered approach combining Ensembl BioMart, g:Profiler, and HomoloGene. Enrichment analysis was conducted separately for each organ and timepoint. The top 10 enriched TFs per condition were visualized using faceted bubble plots. A presence–absence matrix was used to classify TFs as shared or organ-unique and visualized using concentric donut plots and Venn diagrams. TF–target interactions were mapped using ChEA and overlaid with expression heatmaps to highlight regulatory architecture. Target gene sets were further subjected to GO and Hallmark pathway analysis.

To capture global variation in TFs activity, dimensionality reduction was performed on the TFs $\log_2$FC matrix using PCA and UMAP. Organ-specific trajectories were visualized across the time course (D7 to D35). To determine the most organ-discriminative TFs at each timepoint, we trained Random Forest classifiers independently for each day using TFs $\log_2$FC values as predictors and organ identity as the response variable. The Mean decrease in Gini was used to rank TFs importance and highlight top organ-informative regulators.

**Transcriptomic-GWAS integration analysis**

To link transcriptomic signatures with human disease relevance, we integrated rat DEGs with human GWAS loci for hypertension and CKD (NHGRI-EBI GWAS Catalog, release e114_r2025-05-13). Rat–human orthologs were derived as described in earlier section. Overlap statistics were evaluated using Fisher's exact test and overlap genes (hypertension: n = 100; CKD: n = 143) were profiled across all organs–timepoint combinations. Z-score normalized heatmaps were used to assess expression dynamics. Over-representation analysis (GO-BP, KEGG, Reactome) was performed using clusterProfiler and ReactomePA. PPI networks were reconstructed for overlap genes and analyzed using STRING and Louvain clustering to identify disease-relevant modules, which were annotated by GO enrichment and visualized with ggraph.

**Drug perturbation analysis**

To systematically identify compounds that can rescue salt-induced hypertension-associated transcriptional changes, overrepresentation analyses were performed against the Library of Integrated Network-based Cellular Signatures (LINCS) L1000 chemical perturbation databases using enrichr[6]. The LINCS L1000 resource provides a large compendium of gene-expression profiles elicited by hundreds of small molecules across diverse human cell types, thereby enabling direct linkage of disease signatures to candidate pharmacological modulators[7,8]. For each DEGs set, upregulated genes were queried against LINCS_L1000_Chem_Pert_down signatures, representing compounds predicted to reverse the observed gene induction, whereas downregulated genes were tested against LINCS_L1000_Chem_Pert_up signatures, representing compounds predicted to restore suppressed gene expression. The top ten enriched compounds per group were retained. The enrichment results were merged and duplicate entries with identical compound labels collapsed by retaining the instance with the lowest adjusted *P* value. The compound terms that differed in exposure time were also preserved as distinct entries. To provide complementary biological context for the chemical perturbation results, enrichment against the MSigDB Hallmark 2020 gene sets was conducted in parallel. For each gene set, the top five Hallmark pathways were selected based on adjusted *P* values, supporting the mechanistic interpretation of compound–disease relationships by anchoring transcriptional signatures within biological processes. Sankey diagrams were constructed to visualize temporal and pharmacological relationships. Edges connecting each time-point to enriched Hallmark pathways had line widths scaled as $5 \times (-\log_{10}$ adjusted $P$ value) to emphasize temporal signal strength, while edges linking pathways to LINCS compounds were weighted proportionally to the unscaled $-\log_{10}$ adjusted $P$ value. Networks were generated using the *networkD3* R package. Further, identified drugs were categorized manually based on their reported primary targets or mechanisms of action then collapsed into broader therapeutic categories: receptor tyrosine kinase (RTK) inhibitors (including compounds targeting EGFR, HER2, VEGFR, FGFR, ALK, and related kinases), MAPK pathway inhibitors (targeting BRAF, RAF, MEK, ERK, and p38), cell-cycle kinase inhibitors (encompassing CDK, PLK, and CHK inhibitors), PI3K/mTOR inhibitors, HSP90 inhibitors, and anti-inflammatory agents (including JAK, JNK, IKK, NF-κB). Compounds that did not match any of these criteria were classified as "Other." For each tissue and timepoint we created an aggregate enrichment score by summing the $-\log_{10}$ of the adjusted *P* values for every compound in a given drug class. These class-level scores were winsorized at 5000 to prevent extremely small *P* values from dominating the visualization. Streamgraphs were then produced with the *ggstream* and *ggplot2* R packages.

#### Histopathological and immunofluorescence analysis

For histological evaluation of the kidney, liver, and heart, tissue samples were fixed in 10% zinc-buffered formalin, and paraffin-embedded blocks were prepared. Sections of 4 μm thickness were cut and stained with Masson's trichrome for fibrosis assessment and hematoxylin and eosin (H&E) for injury scoring. Whole-slide images were scanned, and quantitative analyses were performed using QuPath software (v0.5.1)[9]. For kidney injury scoring, each whole-kidney section was divided into 1000 × 500 μm fields, and the cortex and medulla were evaluated separately using a semi-quantitative scoring system. Scores were assigned based on the percentage of tissue area affected by necrosis, brush border loss, cast formation, and tubular dilation: 0 = no injury; 1 = <10%; 2 = 11–25%; 3 = 26–50%; and 4 = >50% injury in the field. More than 100 cortical fields and over 50 medullary fields were analyzed per animal. Fibrosis quantification was performed by calculating the percentage of Masson's trichrome positive area separately for cortex and medulla. In the heart and liver, perivascular fibrosis was quantified. Over 20 vessels per sample were selected based on size and shape. Collagen surrounding these vessels was measured as perivascular collagen. The percentage of collagen was calculated by analyzing the ratio of positively stained (blue) pixels to total pixels within each vessel's perivascular region.

#### Serum and urine analysis

Electrolyte and creatinine concentrations in urine and plasma were measured using a blood gas analyzer (ABL800 FLEX, Radiometer America Inc.). Urinary albumin concentrations were quantified using a fluorescence-based dye-binding assay adapted for rat samples. A dilution series of rat serum albumin standards (0.0032–0.2 mg/mL) was prepared in Buffer B (standard diluent: Milli-Q water with buffering salts), and rat urine samples were diluted 1:10 in the same buffer. Twenty-five microliters of each standard or sample were plated in duplicate into a 96-well plate. A working dye solution was prepared fresh by diluting lyophilized AM3 dye (resuspended in isopropanol) 1:50 in Buffer A, a MOPS-based buffer containing 10% isopropanol. Following the addition of 150 μL of working dye per well, plates were incubated for 5 minutes at room temperature in the dark. Fluorescence was measured using a microplate reader (excitation: 560 nm; emission: 620 nm). Albumin concentrations were calculated by interpolating sample values from a standard, corrected for blank values, and dilution factor as reported previously[1].

#### Statistics

Statistical analyses were performed using GraphPad Prism (v10.5) and R (v4.4.0). For RNA-seq data, differential gene expressions were assessed using DESeq2. Genes with an $|\log_2 \text{fold change}| \geq 0.585$, equivalent to a fold change ≥ 1.5, adjusted $P < 0.05$ were considered significantly differentially expressed. Statistical significance was assessed using one-way or two-way ANOVA followed by Šidák's multiple comparisons test for multiple groups. Data are presented as mean ± standard error of the mean (SEM), and $P < 0.05$ was considered statistically significant.

**Study approval**

All animal procedures were conducted in accordance with the Guide for the Care and Use of Laboratory Animals (National Academies Press, 2011) and were approved by the Institutional Animal Care and Use Committee (IACUC) at the University of South Florida.

#### SUPPLEMENTAL FIGURES

**Figure S1**. **Effects of high salt diet on blood pressure and tissue-specific transcriptional profiles**. **A**. Mean arterial pressure (MAP) in male Dahl SS rats fed normal salt (NS) or high salt (HS, 4% NaCl) diet. Two-way repeated-measures ANOVA was used. The inset box summarizes the two-way ANOVA results, showing significant main effects of time (*****P* < 0.0001) and diet (***P* < 0.01), as well as a significant time and diet interaction (*****P* < 0.0001). Post hoc pairwise comparisons between HS and NS at each time point were performed with Sidak's correction and significance levels are indicated on the graph as **P* < 0.05, ***P* < 0.01, ****P* < 0.001, and *****P* < 0.0001, with ns indicating not significant. Data are presented as mean ± standard error of the mean (SEM). **B**. Principal component analysis (PCA) of variance-stabilized (VST) RNA-seq counts from kidney cortex, kidney medulla, liver, and heart tissues obtained from a separate cohort (not subjected to telemetry surgery to avoid surgery-related transcriptomic changes) under NS conditions and after 7, 14, 21, and 35 days of HS diet. **C**. Rank–abundance distributions of transcript counts [$\log_2$(CPM + 1)] for each organ across experimental time points. Genes are ordered from highest to lowest mean expression on the x-axis, with abundance plotted on the y-axis. Colored lines show consistent global transcript abundance distributions across organs and time points. CX, cortex; MD, medulla; LV, liver; HR, heart; D7, day 7; D14, day 14; D21, day 21; and D35, day 35 time points. *n* = 6 male rats per group.

**Figure S2**. **Volcano plot analysis reveals organ-specific and temporal patterns of gene regulation under high salt diet-induced hypertension**. Volcano plots showing differential gene expression ($\log_2$ fold change vs $-\log_{10}$ adjusted *P* value) in the kidney cortex, kidney medulla, liver, and heart at days 7, 14, 21, and 35 relatives to their respective controls. Significantly upregulated genes are shown in red, downregulated genes in blue, and non-significant genes in gray. Numeric labels above each plot indicate counts of upregulated (Up), downregulated (Down), and non-significant (NS) genes. Genes with $|\log_2 \text{fold change}| \geq 0.585$, equivalent to a fold change ≥ 1.5 with adjusted *P* < 0.05 were considered significant. CX, kidney cortex; MD, kidney medulla; LV, liver; HR, heart; D7, day 7; D14, day 14; D21, day 21; and D35, day 35 time points. adj. p indicated adjusted *P* value. *n* = 6 male rats per group.

**Figure S3**. **Electrolyte, histological and molecular changes during salt-induced hypertension in Dahl SS rats**. **A–E**. Electrolyte measurements showing urinary sodium-to-creatinine ratio (Na/Cr) (**A**), urinary chloride-to-creatinine ratio (Cl/Cr) (**B**), blood sodium (Na) (**C**), blood chloride (Cl) (**D**), and blood pH (**E**) in normal-salt controls (NS) and at days 7, 14, 21, and 35 of high salt diet. **F**. Representative Masson's trichrome images of liver and heart from NS and days 7, 14, 21, and 35 after high salt diet groups. Collagen (fibrosis) is blue; parenchyma/myocardium is red. Scale bars, 100 µm. **G**. Semi-quantitative analysis of Trichrome-positive area (%) in liver and heart (perivascular region) across time points. **H**. Heatmaps are showing $\log_2$ fold-change (vs NS) expression of selected genes associated with fibrosis and inflammation at each time point in liver and heart. Expression levels of genes was extracted from RNA-seq analysis. For **A–G**, Data are shown mean ± standard error of the mean (SEM). One-way ANOVA with Šídák's multiple comparisons test was used. Statistical comparison is vs NS; ns, not significant; **P* < 0.05; ***P* < 0.01; *****P* < 0.0001; n = 4–8 male rats per group. For **H**, Asterisks denote Benjamini–Hochberg adjusted-*P* thresholds (*adj. P*). **adj. P* < 0.05; ***adj. P < 0.01;* ****adj. P* < 0.001; *n* = 6 male rats per group. D7, day 7; D14, day 14; D21, day 21; and D35, day 35 time points. *Vcam1*, v*ascular cell adhesion molecule 1*; *Tgfb1*, *transforming growth factor beta 1*; *Icam1*, *intercellular adhesion molecule 1*; *Got1*, *glutamic-oxaloacetic transaminase 1*; *Col3a1*, c*ollagen type III alpha-1 chain*; *Loxl2, lysyl oxidase-like 2*.

**Figure S4**. **Temporal dynamics of highly variable pathways across organs**. Line plots showing temporal trajectories of the top 10 most variable Hallmark pathways within each tissue. Values represent the mean $\log_2$ fold change ($\log_2$FC) relative to controls at each time point. The dotted line denotes control levels; pathways above the line are upregulated, while those below are downregulated. CX, kidney cortex; MD, kidney medulla; LV, liver; HR, heart; D7, day 7; D14, day 14; D21, day 21; and D35, day 35 time points. $n = 6$ male rats per group.

**Figure S5**. **Unsupervised clustering of hallmark pathway trajectories**. **A**. UMAP visualization of Hallmark pathway profiles shows four distinct clusters based on their expression patterns across different tissues and time points (days 7–35). Each point represents a pathway, colored by its assigned cluster. Pathways in Cluster 1 (red) are enriched for mitochondrial and lipid metabolism processes. Cluster 2 (blue) includes immune and inflammatory response pathways. Cluster 3 (green) contains a broad range of metabolic, stress response, and developmental pathways. Cluster 4 (purple) groups cell cycle–related pathways including E2F targets and G2M checkpoint. **B**. Silhouette score plot used to determine the optimal number of clusters (k = 4), based on average silhouette width across k = 2–8. EMT, epithelial–mesenchymal transition. $n = 6$ male rats per group.

**Figure S6**. **GO Biological Process (GO-BP) enrichment of genes commonly dysregulated across all four tissues in high salt diet-induced hypertension**. Differentially expressed genes shared among the kidney cortex, kidney medulla, liver, and heart at days 7 and 35 were analyzed for Gene Ontology Biological Process (GO-BP) enrichment. **A** and **B** show results for day 7, with panel **A** depicting the top 10 enriched GO-BP terms and panel **B** showing the GO-BP enrichment map visualizing semantic similarity networks among enriched terms. **C** and **D** show results for day 35, with panel **C** depicting the top 10 enriched GO-BP terms and panel **D** showing the GO-BP enrichment map visualizing semantic similarity networks among enriched terms. Node size reflects gene set size and color intensity indicates statistical significance. D7, day 7; D35, day 35; adj.p, adjusted *P* value. $n = 6$ male rats per group.

**Figure S7**. **High salt diet-induced hypertension dynamically regulates biological processes across organs and time points**. Bubble plots show the top 5 enriched GO Biological Process (GO-BP) terms (adjusted $P < 0.05$) identified from all significant differentially expressed genes ($|\log_2$ fold change$| \geq 0.585$, equivalent to a fold change $\geq 1.5$ with adjusted $P < 0.05$) in the kidney cortex, kidney medulla, liver, and heart at days 7, 14, 21, and 35. Dot size represents the number of overlapping genes, and color denotes statistical significance ($-\log_{10}$ adjusted $P$). Results highlight dynamic organ- and time-specific regulation of processes including cell division, immune response, metabolic activity, and tissue-specific programs. CX, cortex; MD, medulla; LV, liver; HR, heart; D7, day 7; D14, day 14; D21, day 21; and D35, day 35 time points. adj. p indicated adjusted *P* value. *n* = 6 male rats per group.

**Figure S8**. **Distinct temporal and organ-specific Hallmark pathways engaged by high salt diet-induced hypertension**. Bubble plots depict the top enriched Hallmark pathways (adjusted $P < 0.05$) derived from significantly altered genes ($|\log_2$ fold change$| \geq 0.585$, equivalent to a fold change $\geq 1.5$ with adjusted $P < 0.05$) in the kidney cortex, kidney medulla, liver, and heart at days 7, 14, 21, and 35. Circle size indicates the number of overlapping genes, and color denotes statistical significance ($-\log_{10}$ adjusted *P*). CX, cortex; MD, medulla; LV, liver; HR, heart; D7, day 7; D14, day 14; D21, day 21; and D35, day 35 time points. adj. p indicated adjusted *P* value. *n* = 6 male rats per group.

**Figure S9**. **Transcription factor enrichment analysis using ChIP-X Enrichment Analysis (ChEA) reveals dynamic, tissue- and time-specific regulators of high salt diet-induced hypertension**. Bubble plots show the top enriched transcription factors identified by ChEA analysis (adjusted $P < 0.05$) using significantly altered genes from the kidney cortex, kidney medulla, liver, and heart at days 7, 14, 21, and 35. Dot size represents the number of overlapping target genes regulated by each transcription factor, while color intensity denotes statistical significance ($-\log_{10}$ adjusted *P*). Results highlight distinct temporal and tissue-specific TFs underlying hypertension-associated transcriptional remodeling. TF, transcription factors; CX, cortex; MD, medulla; LV, liver; HR, heart; D7, day 7; D14, day 14; D21, day 21; and D35, day 35 time points. adj. p indicated adjusted *P* value. *n* = 6 male rats per group.

**Figure S10**. **Temporal dynamics and organ-specific divergence of transcription factor programs in response to high salt diet–induced hypertension. A**. Heatmap showing $\log_2$-fold changes of 79 transcription factors differentially expressed ($|\log_2$ fold change$| \geq 0.585$, equivalent to a fold change $\geq 1.5$; adjusted $P < 0.05$) across the kidney cortex, kidney medulla, liver, and heart. **B**. Principal Component Analysis (PCA, top) and Uniform Manifold Approximation and Projection (UMAP, bottom) of transcription factor $\log_2$-fold-change profiles, capturing tissue and temporal trajectories from day 7 to day 35. Time points were connected sequentially to guide visualization of transcriptional shifts across the disease course within each tissue. Divergent transcription factor program evolution is most evident in the medulla and liver, with subtler differences in the heart and cortex. **C**. Top ten transcription factors most discriminatory for organ type at each time point, identified by Random Forest classification. Bar lengths indicate variable importance, measured by mean decrease in Gini, reflecting dynamic shifts in transcriptional regulators that drive organ-specific responses over time. TF, transcription factors; CX, cortex; MD, medulla; LV, liver; HR, heart; D7, day 7; D14, day 14; D21, day 21; and D35, day 35 time points. *n* = 6 male rats per group.

**Figure S11**. **Temporal and organ-specific dynamics of genes overlapping human GWAS loci for hypertension and chronic kidney disease in salt-induced hypertension**. **A** and **B**. Heatmaps showing Z-scored $\log_2$ fold-changes of rat orthologs overlapping with human GWAS loci for hypertension (**A**) and chronic kidney disease (CKD; **B**) across the kidney cortex, kidney medulla, liver, and heart at days 7, 14, 21, and 35 following high salt diet exposure in Dahl SS rats. CX, cortex; MD, medulla; LV, liver; HR, heart; D7, day 7; D14, day 14; D21, day 21; and D35, day 35 time points. *n* = 6 male rats per group.

**Figure S12**. **GWAS-overlap genes reveal divergent biological pathways for hypertension and chronic kidney disease in Dahl SS rats**. **A**. Comparative Gene Ontology Biological Process (GO-BP) enrichment reveals disease-specific biological processes enriched in the hypertension and CKD overlap sets. **B** and **C**. KEGG and Reactome enriched terms for hypertension-overlap genes. Top pathways include cortisol and renin synthesis and secretion, ECM–receptor interaction, and extracellular matrix organization, implicating stress-responsive and structural remodeling pathways in hypertensive injury. **D** and **E**. KEGG and Reactome enriched terms of CKD-overlap genes reveal enrichment of metabolic processes, including fatty acid oxidation, cytochrome P450-mediated detoxification, bile acid transport, and tryptophan metabolism. adj. p indicated adjusted *P* value.

**Figure S13**. **Temporal and organ-specific prediction of small molecules targeting downregulated gene programs in salt-sensitive hypertension**. **A–D**. Sankey diagrams integrating differential expression profiles with LINCS L1000 chemical perturbation signatures for significantly downregulated genes in the kidney cortex (**A**), kidney medulla (**B**), liver (**C**), and heart (**D**). Each network connects time points (D7–D35; left), enriched top five Hallmark pathways (middle), and the top predicted small molecules (right) prioritized for their potential to restore suppressed biological programs. Edge thickness denotes $-\log_{10}$(adjusted *P* value), indicating enrichment strength. CX, cortex; MD, medulla; LV, liver; HR, heart; D7, day 7; D14, day 14; D21, day 21; and D35, day 35 time points. *n* = 6 male rats per group.

**Figure S1**

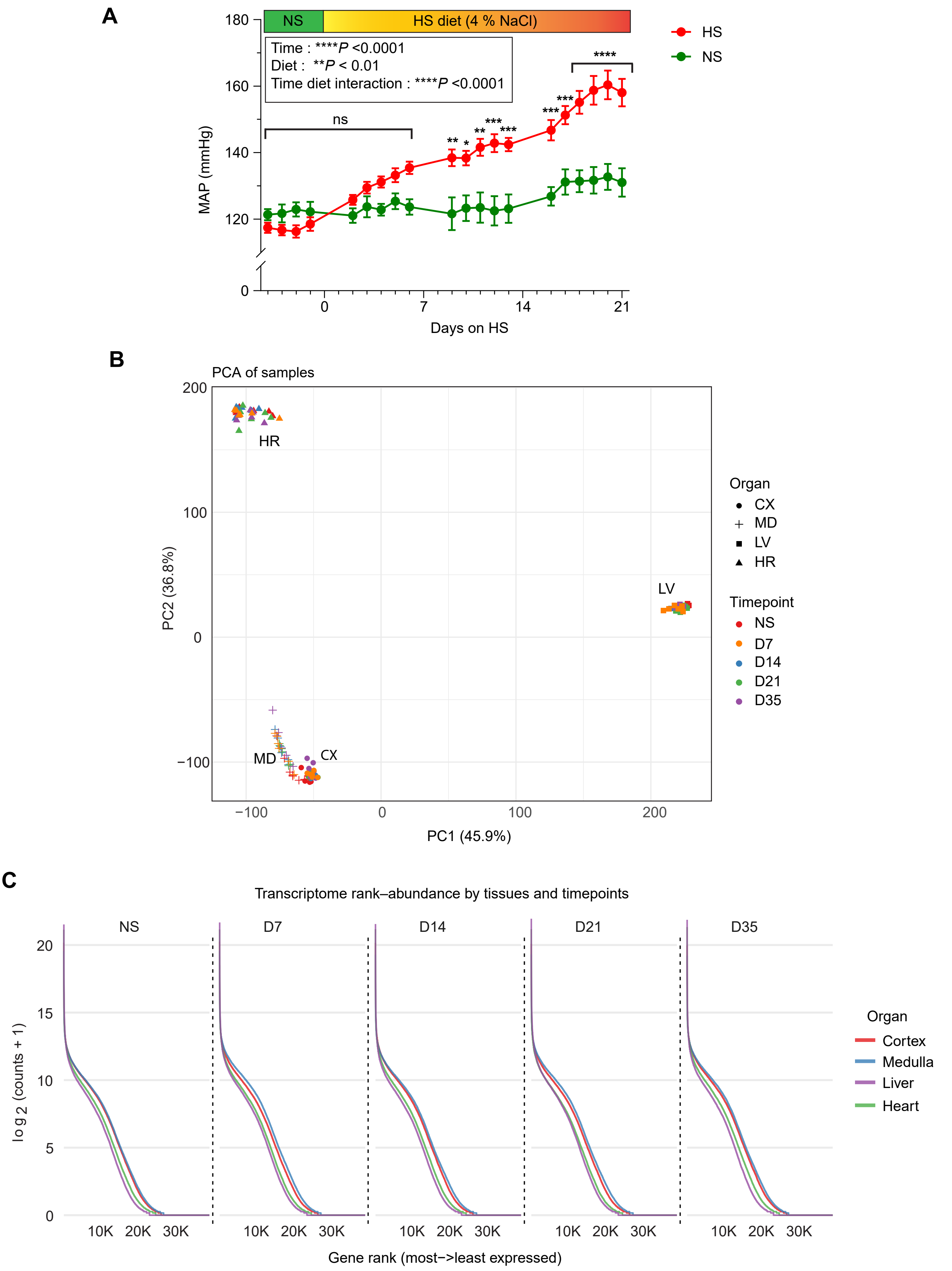
A
NS
HS diet (4 % NaCl)
Time : ****P <0.0001
Diet : **P < 0.01
Time diet interaction : ****P <0.0001
ns
HS
NS
MAP (mmHg)
Days on HS
B
PCA of samples
HR
LV
MD
CX
PC2 (36.8%)
PC1 (45.9%)
Organ
CX
MD
LV
HR
Timepoint
NS
D7
D14
D21
D35
C
Transcriptome rank–abundance by tissues and timepoints
NS
D7
D14
D21
D35
log2 (counts + 1)
Gene rank (most−>least expressed)
Organ
Cortex
Medulla
Liver
Heart

**Figure S2**

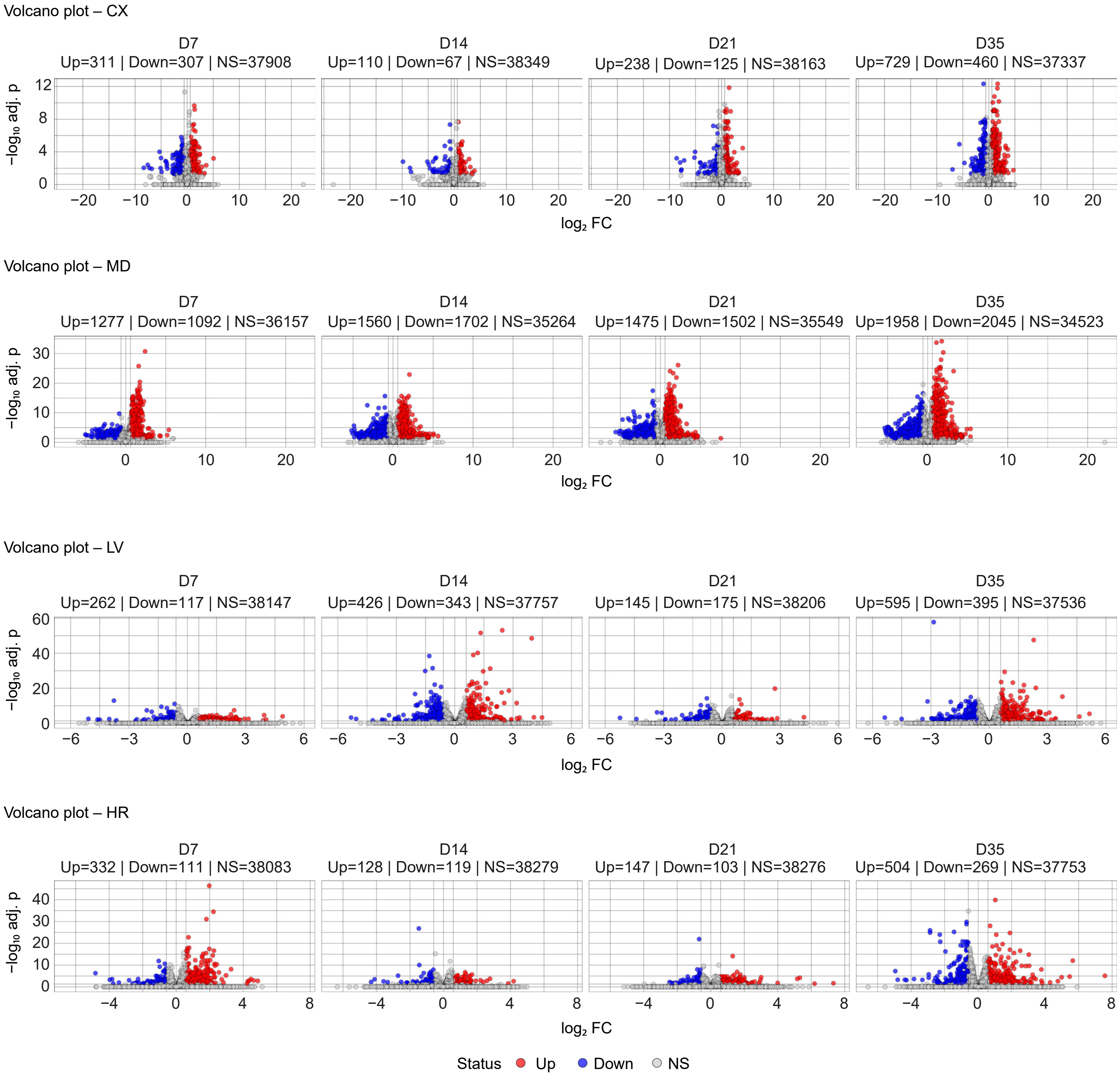
Volcano plot – CX
D7
Up=311 | Down=307 | NS=37908
D14
Up=110 | Down=67 | NS=38349
D21
Up=238 | Down=125 | NS=38163
D35
Up=729 | Down=460 | NS=37337
−log10 adj. p
log2 FC
Volcano plot – MD
D7
Up=1277 | Down=1092 | NS=36157
D14
Up=1560 | Down=1702 | NS=35264
D21
Up=1475 | Down=1502 | NS=35549
D35
Up=1958 | Down=2045 | NS=34523
−log10 adj. p
log2 FC
Volcano plot – LV
D7
Up=262 | Down=117 | NS=38147
D14
Up=426 | Down=343 | NS=37757
D21
Up=145 | Down=175 | NS=38206
D35
Up=595 | Down=395 | NS=37536
−log10 adj. p
log2 FC
Volcano plot – HR
D7
Up=332 | Down=111 | NS=38083
D14
Up=128 | Down=119 | NS=38279
D21
Up=147 | Down=103 | NS=38276
D35
Up=504 | Down=269 | NS=37753
−log10 adj. p
log2 FC
Status Up Down NS

### Figure S3

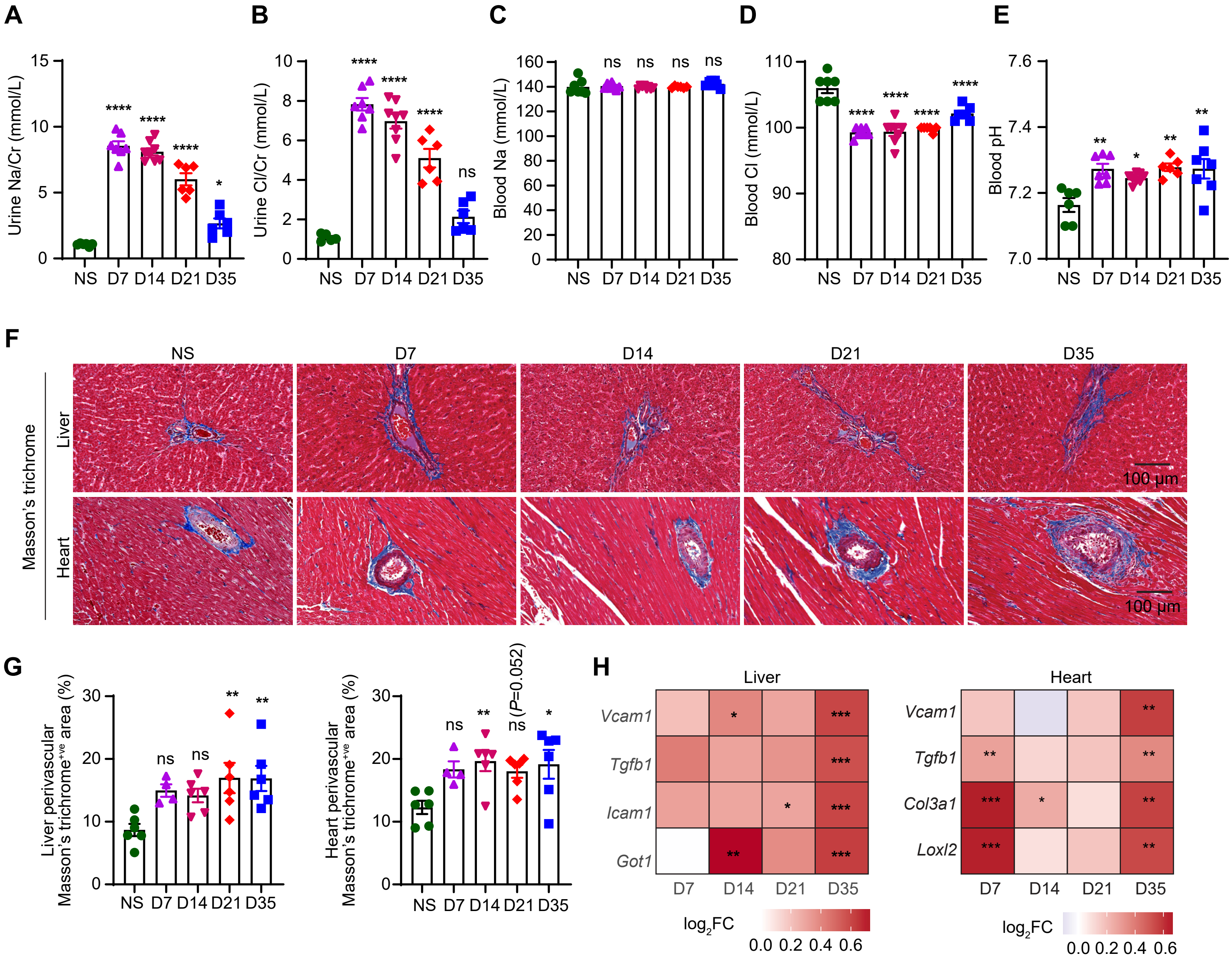
A
B
C
D
E
Urine Na/Cr (mmol/L)
Urine Cl/Cr (mmol/L)
Blood Na (mmol/L)
Blood Cl (mmol/L)
Blood pH
NS D7 D14 D21 D35
F
NS
D7
D14
D21
D35
Masson's trichrome
Liver
Heart
100 µm
G
Liver perivascular Masson's trichrome$^{+ve}$ area (%)
Heart perivascular Masson's trichrome$^{+ve}$ area (%)
(P=0.052)
H
Liver
Heart
Vcam1
Tgfb1
Icam1
Got1
Col3a1
Loxl2
D7 D14 D21 D35
$log_2FC$
0.0 0.2 0.4 0.6

**Figure S4**

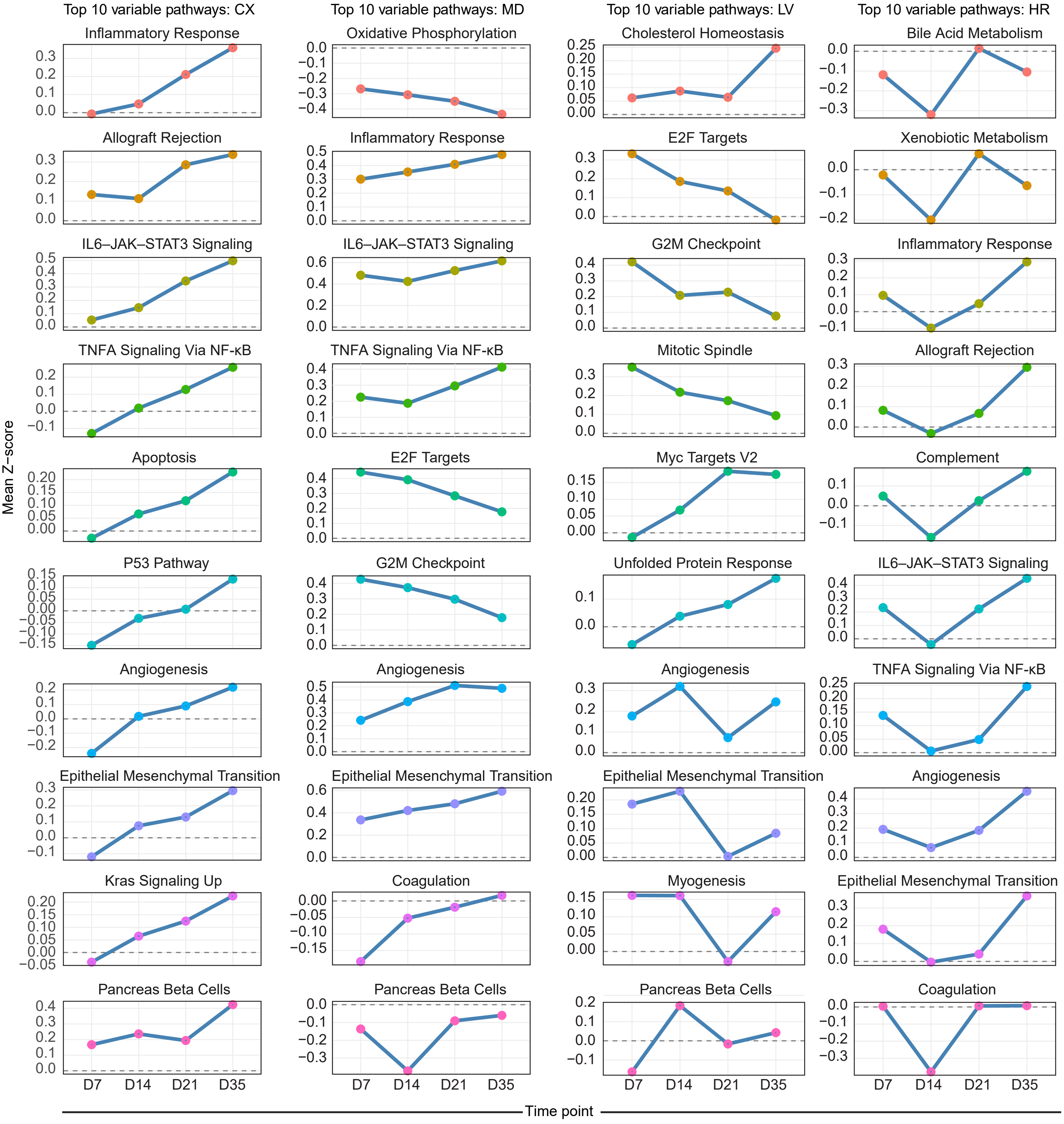

Top 10 variable pathways: CX
Top 10 variable pathways: MD
Top 10 variable pathways: LV
Top 10 variable pathways: HR
Inflammatory Response
Oxidative Phosphorylation
Cholesterol Homeostasis
Bile Acid Metabolism
Allograft Rejection
Inflammatory Response
E2F Targets
Xenobiotic Metabolism
IL6–JAK–STAT3 Signaling
IL6–JAK–STAT3 Signaling
G2M Checkpoint
Inflammatory Response
TNFA Signaling Via NF-κB
TNFA Signaling Via NF-κB
Mitotic Spindle
Allograft Rejection
Apoptosis
E2F Targets
Myc Targets V2
Complement
P53 Pathway
G2M Checkpoint
Unfolded Protein Response
IL6–JAK–STAT3 Signaling
Angiogenesis
Angiogenesis
Angiogenesis
TNFA Signaling Via NF-κB
Epithelial Mesenchymal Transition
Epithelial Mesenchymal Transition
Epithelial Mesenchymal Transition
Angiogenesis
Kras Signaling Up
Coagulation
Myogenesis
Epithelial Mesenchymal Transition
Pancreas Beta Cells
Pancreas Beta Cells
Pancreas Beta Cells
Coagulation
Mean Z–score
D7
D14
D21
D35
Time point

**Figure S5**

**A**

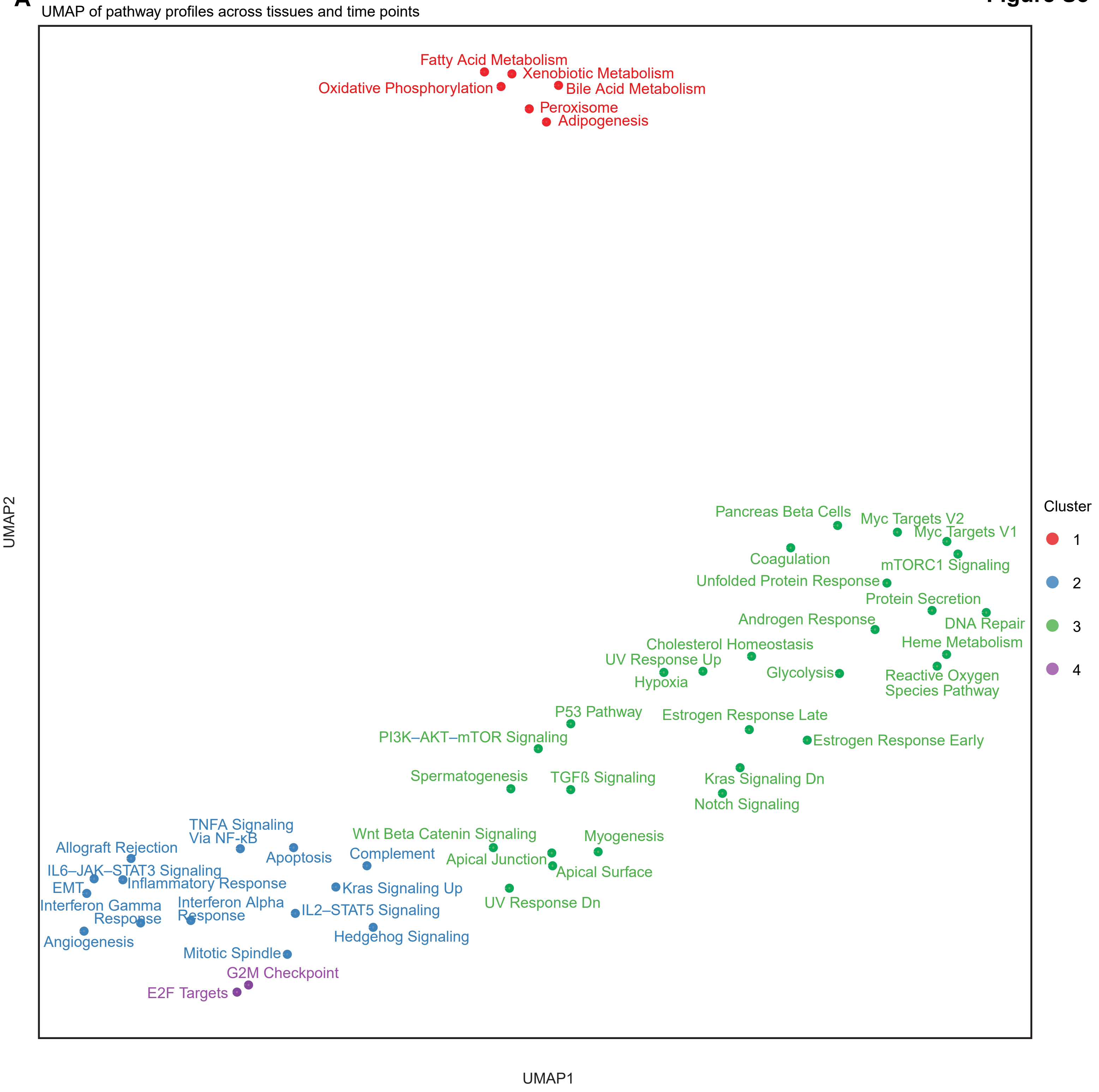
UMAP of pathway profiles across tissues and time points
Fatty Acid Metabolism
Xenobiotic Metabolism
Oxidative Phosphorylation
Bile Acid Metabolism
Peroxisome
Adipogenesis
Pancreas Beta Cells
Myc Targets V2
Myc Targets V1
Coagulation
mTORC1 Signaling
Unfolded Protein Response
Protein Secretion
Androgen Response
DNA Repair
Cholesterol Homeostasis
Heme Metabolism
UV Response Up
Glycolysis
Hypoxia
Reactive Oxygen Species Pathway
P53 Pathway
Estrogen Response Late
PI3K–AKT–mTOR Signaling
Estrogen Response Early
Spermatogenesis
TGFß Signaling
Kras Signaling Dn
Notch Signaling
TNFA Signaling Via NF-κB
Allograft Rejection
Wnt Beta Catenin Signaling
Myogenesis
Apoptosis
Complement
Apical Junction
IL6–JAK–STAT3 Signaling
Apical Surface
Inflammatory Response
EMT
Kras Signaling Up
Interferon Gamma Response
Interferon Alpha Response
UV Response Dn
IL2–STAT5 Signaling
Angiogenesis
Hedgehog Signaling
Mitotic Spindle
G2M Checkpoint
E2F Targets
UMAP1
UMAP2
Cluster
1
2
3
4

**B**

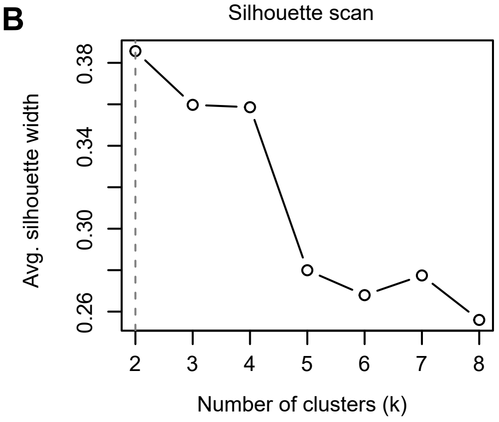
Silhouette scan
Avg. silhouette width
0.26
0.30
0.34
0.38
Number of clusters (k)
2
3
4
5
6
7
8

**Figure S6**

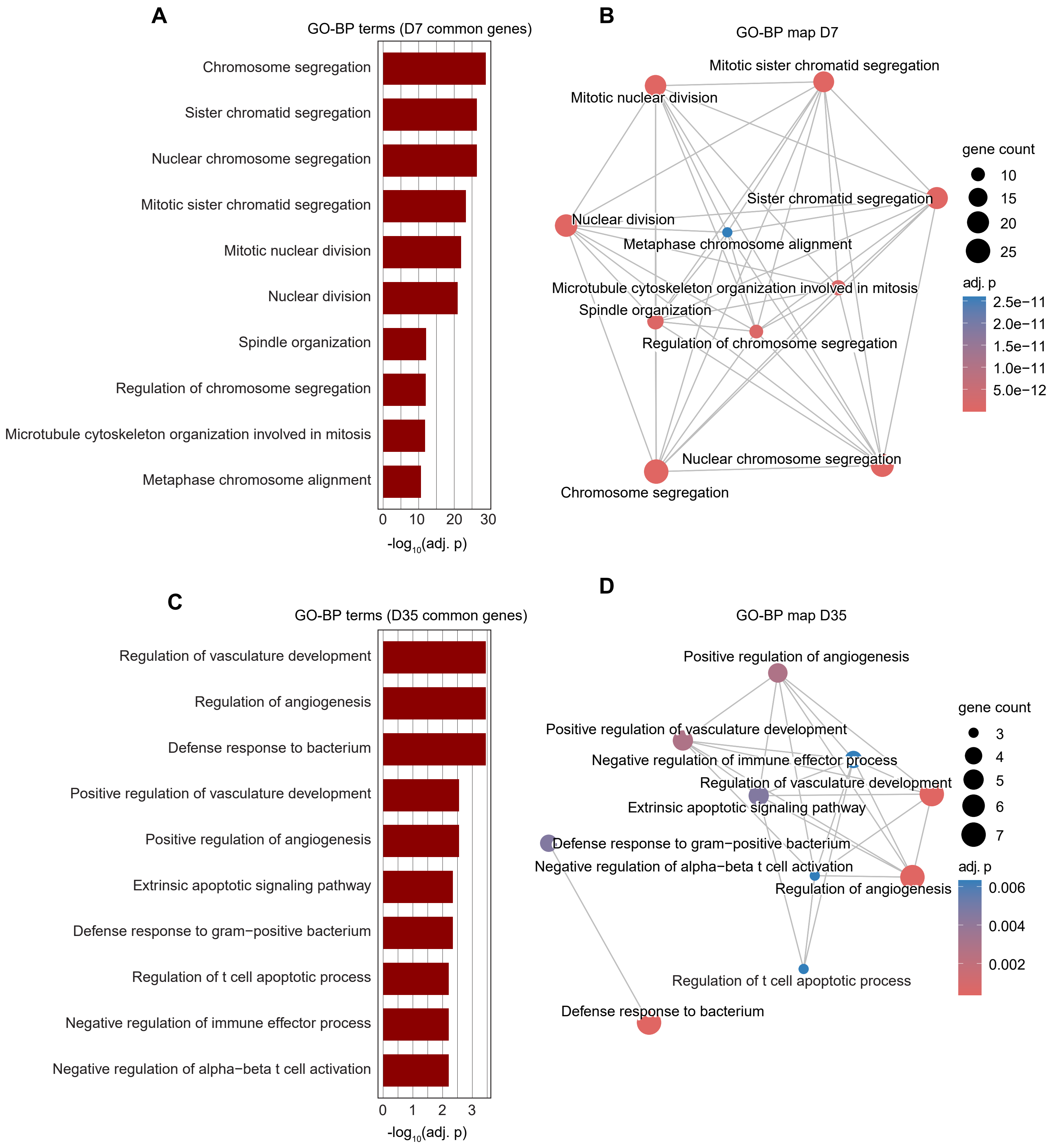

A
GO-BP terms (D7 common genes)
Chromosome segregation
Sister chromatid segregation
Nuclear chromosome segregation
Mitotic sister chromatid segregation
Mitotic nuclear division
Nuclear division
Spindle organization
Regulation of chromosome segregation
Microtubule cytoskeleton organization involved in mitosis
Metaphase chromosome alignment
0 10 20 30
-log10(adj. p)
B
GO-BP map D7
Mitotic sister chromatid segregation
Mitotic nuclear division
Sister chromatid segregation
Nuclear division
Metaphase chromosome alignment
Microtubule cytoskeleton organization involved in mitosis
Spindle organization
Regulation of chromosome segregation
Nuclear chromosome segregation
Chromosome segregation
gene count
10
15
20
25
adj. p
2.5e−11
2.0e−11
1.5e−11
1.0e−11
5.0e−12
C
GO-BP terms (D35 common genes)
Regulation of vasculature development
Regulation of angiogenesis
Defense response to bacterium
Positive regulation of vasculature development
Positive regulation of angiogenesis
Extrinsic apoptotic signaling pathway
Defense response to gram−positive bacterium
Regulation of t cell apoptotic process
Negative regulation of immune effector process
Negative regulation of alpha−beta t cell activation
0 1 2 3
-log10(adj. p)
D
GO-BP map D35
Positive regulation of angiogenesis
Positive regulation of vasculature development
Negative regulation of immune effector process
Regulation of vasculature development
Extrinsic apoptotic signaling pathway
Defense response to gram−positive bacterium
Negative regulation of alpha−beta t cell activation
Regulation of angiogenesis
Regulation of t cell apoptotic process
Defense response to bacterium
gene count
3
4
5
6
7
adj. p
0.006
0.004
0.002

### Figure S7

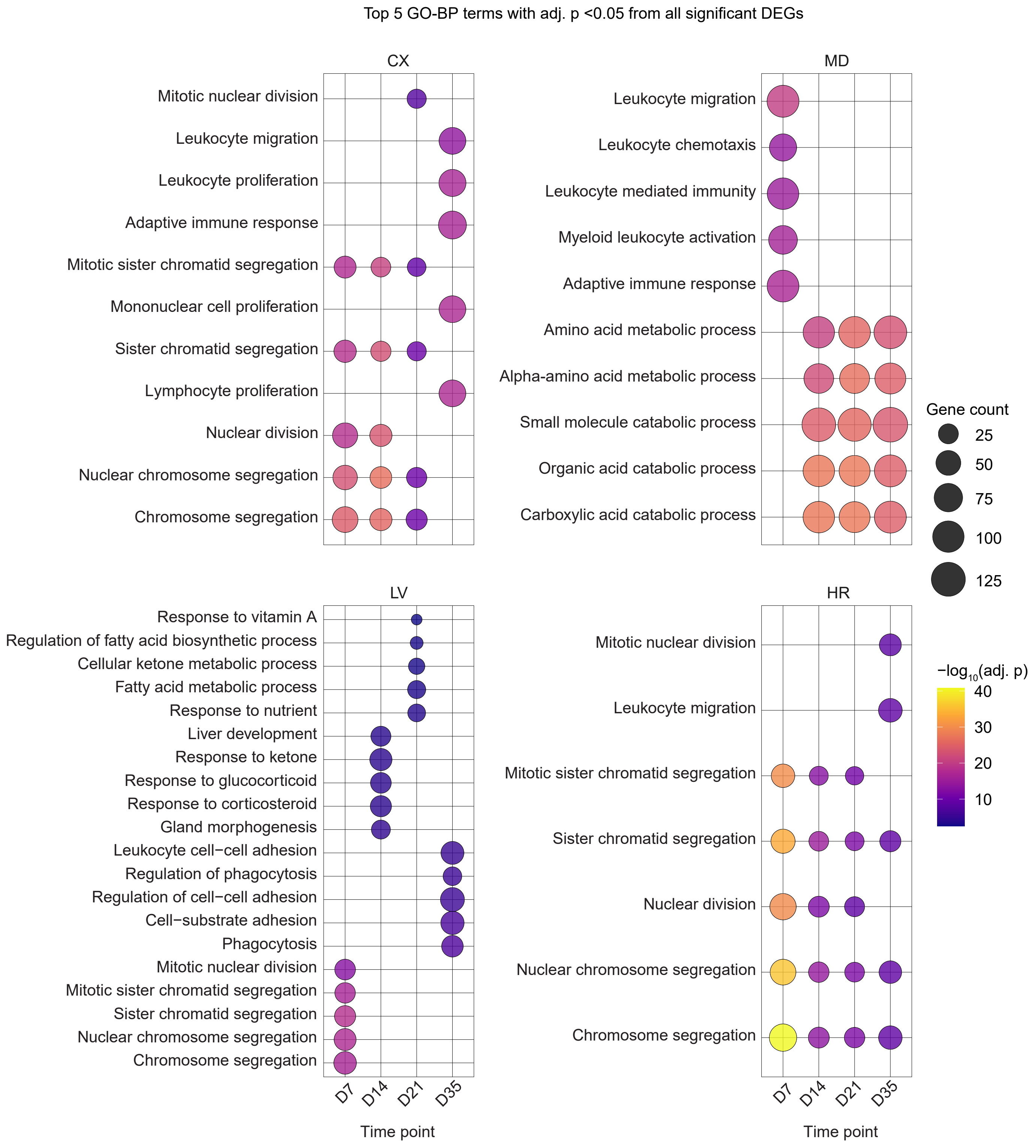

Top 5 GO-BP terms with adj. p <0.05 from all significant DEGs
CX
Mitotic nuclear division
Leukocyte migration
Leukocyte proliferation
Adaptive immune response
Mitotic sister chromatid segregation
Mononuclear cell proliferation
Sister chromatid segregation
Lymphocyte proliferation
Nuclear division
Nuclear chromosome segregation
Chromosome segregation
MD
Leukocyte migration
Leukocyte chemotaxis
Leukocyte mediated immunity
Myeloid leukocyte activation
Adaptive immune response
Amino acid metabolic process
Alpha-amino acid metabolic process
Small molecule catabolic process
Organic acid catabolic process
Carboxylic acid catabolic process
LV
Response to vitamin A
Regulation of fatty acid biosynthetic process
Cellular ketone metabolic process
Fatty acid metabolic process
Response to nutrient
Liver development
Response to ketone
Response to glucocorticoid
Response to corticosteroid
Gland morphogenesis
Leukocyte cell−cell adhesion
Regulation of phagocytosis
Regulation of cell−cell adhesion
Cell−substrate adhesion
Phagocytosis
Mitotic nuclear division
Mitotic sister chromatid segregation
Sister chromatid segregation
Nuclear chromosome segregation
Chromosome segregation
HR
Mitotic nuclear division
Leukocyte migration
Mitotic sister chromatid segregation
Sister chromatid segregation
Nuclear division
Nuclear chromosome segregation
Chromosome segregation
D7
D14
D21
D35
Time point
Gene count
25
50
75
100
125
−log10(adj. p)
40
30
20
10

**Figure S8**

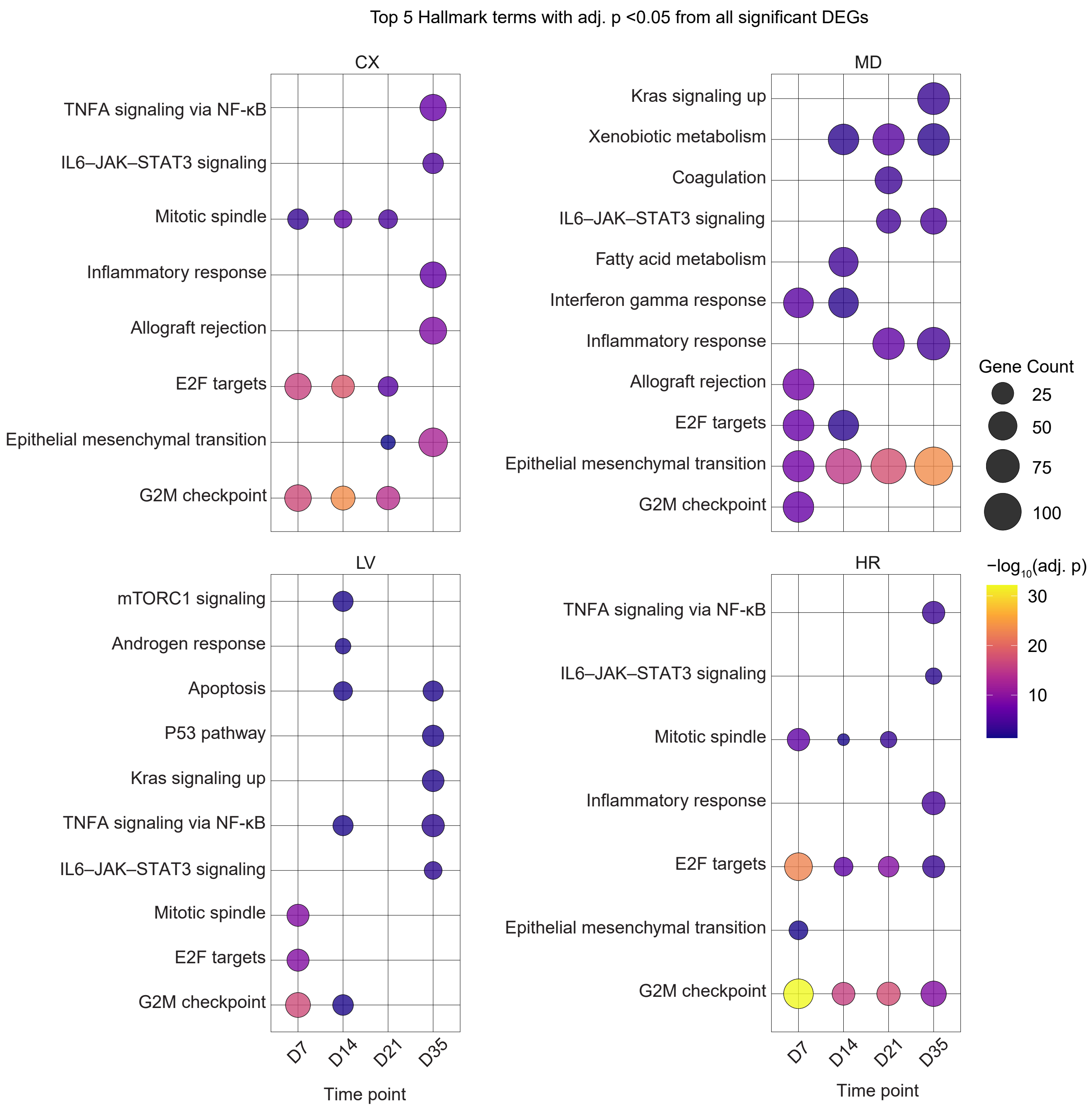
Top 5 Hallmark terms with adj. p <0.05 from all significant DEGs
CX
TNFA signaling via NF-κB
IL6–JAK–STAT3 signaling
Mitotic spindle
Inflammatory response
Allograft rejection
E2F targets
Epithelial mesenchymal transition
G2M checkpoint
MD
Kras signaling up
Xenobiotic metabolism
Coagulation
IL6–JAK–STAT3 signaling
Fatty acid metabolism
Interferon gamma response
Inflammatory response
Allograft rejection
E2F targets
Epithelial mesenchymal transition
G2M checkpoint
LV
mTORC1 signaling
Androgen response
Apoptosis
P53 pathway
Kras signaling up
TNFA signaling via NF-κB
IL6–JAK–STAT3 signaling
Mitotic spindle
E2F targets
G2M checkpoint
HR
TNFA signaling via NF-κB
IL6–JAK–STAT3 signaling
Mitotic spindle
Inflammatory response
E2F targets
Epithelial mesenchymal transition
G2M checkpoint
D7
D14
D21
D35
Time point
Gene Count
25
50
75
100
−log10(adj. p)
30
20
10

### Figure S9

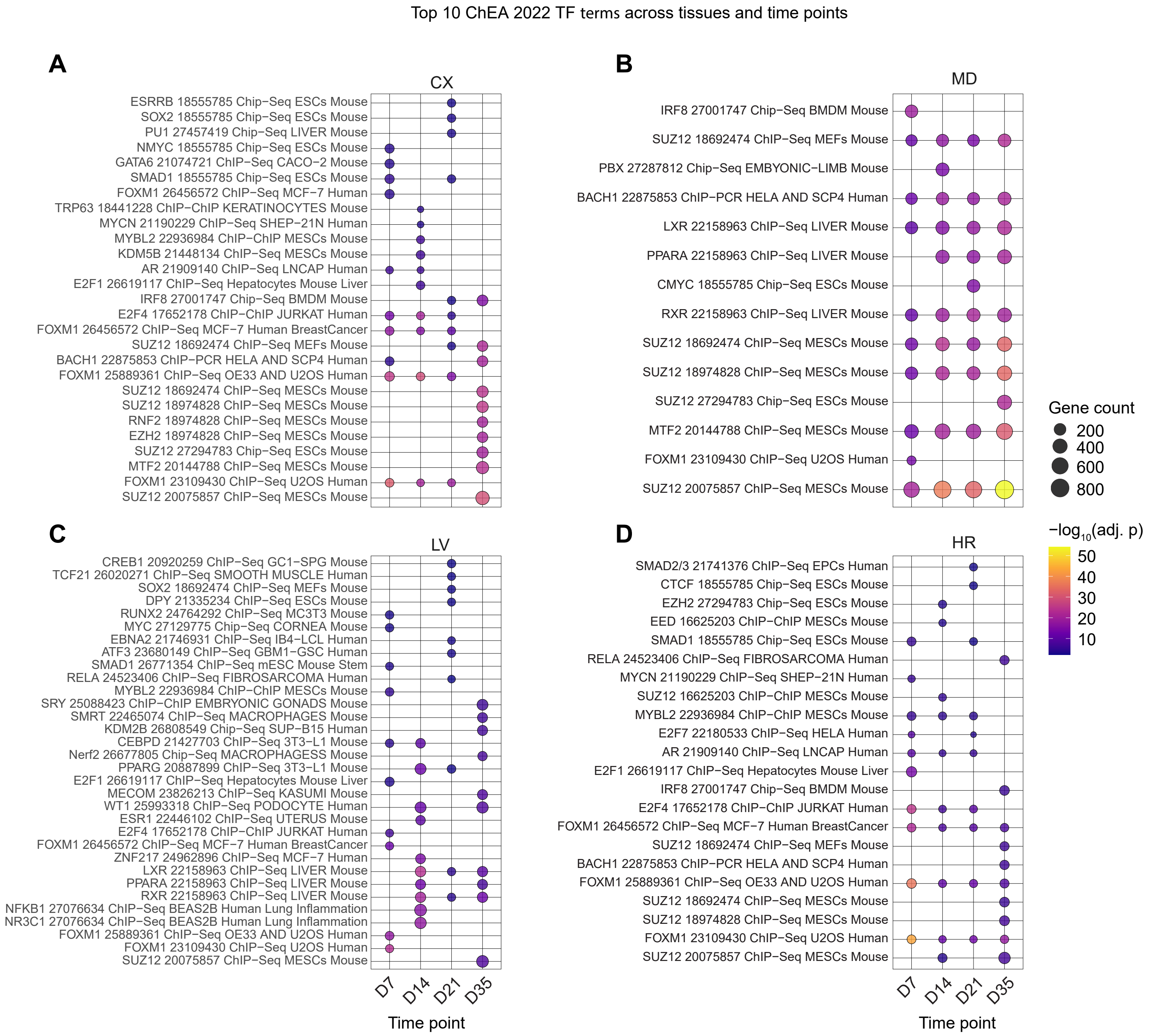
Top 10 ChEA 2022 TF terms across tissues and time points
A
CX
ESRRB 18555785 Chip−Seq ESCs Mouse
SOX2 18555785 Chip−Seq ESCs Mouse
PU1 27457419 Chip−Seq LIVER Mouse
NMYC 18555785 Chip−Seq ESCs Mouse
GATA6 21074721 ChIP−Seq CACO−2 Mouse
SMAD1 18555785 Chip−Seq ESCs Mouse
FOXM1 26456572 ChIP−Seq MCF−7 Human
TRP63 18441228 ChIP−ChIP KERATINOCYTES Mouse
MYCN 21190229 ChIP−Seq SHEP−21N Human
MYBL2 22936984 ChIP−ChIP MESCs Mouse
KDM5B 21448134 ChIP−Seq MESCs Mouse
AR 21909140 ChIP−Seq LNCAP Human
E2F1 26619117 ChIP−Seq Hepatocytes Mouse Liver
IRF8 27001747 Chip−Seq BMDM Mouse
E2F4 17652178 ChIP−ChIP JURKAT Human
FOXM1 26456572 ChIP−Seq MCF−7 Human BreastCancer
SUZ12 18692474 ChIP−Seq MEFs Mouse
BACH1 22875853 ChIP−PCR HELA AND SCP4 Human
FOXM1 25889361 ChIP−Seq OE33 AND U2OS Human
SUZ12 18692474 ChIP−Seq MESCs Mouse
SUZ12 18974828 ChIP−Seq MESCs Mouse
RNF2 18974828 ChIP−Seq MESCs Mouse
EZH2 18974828 ChIP−Seq MESCs Mouse
SUZ12 27294783 Chip−Seq ESCs Mouse
MTF2 20144788 ChIP−Seq MESCs Mouse
FOXM1 23109430 ChIP−Seq U2OS Human
SUZ12 20075857 ChIP−Seq MESCs Mouse
B
MD
IRF8 27001747 Chip−Seq BMDM Mouse
SUZ12 18692474 ChIP−Seq MEFs Mouse
PBX 27287812 Chip−Seq EMBYONIC−LIMB Mouse
BACH1 22875853 ChIP−PCR HELA AND SCP4 Human
LXR 22158963 ChIP−Seq LIVER Mouse
PPARA 22158963 ChIP−Seq LIVER Mouse
CMYC 18555785 Chip−Seq ESCs Mouse
RXR 22158963 ChIP−Seq LIVER Mouse
SUZ12 18692474 ChIP−Seq MESCs Mouse
SUZ12 18974828 ChIP−Seq MESCs Mouse
SUZ12 27294783 Chip−Seq ESCs Mouse
MTF2 20144788 ChIP−Seq MESCs Mouse
FOXM1 23109430 ChIP−Seq U2OS Human
SUZ12 20075857 ChIP−Seq MESCs Mouse
Gene count
200
400
600
800
C
LV
CREB1 20920259 ChIP−Seq GC1−SPG Mouse
TCF21 26020271 ChIP−Seq SMOOTH MUSCLE Human
SOX2 18692474 ChIP−Seq MEFs Mouse
DPY 21335234 ChIP−Seq ESCs Mouse
RUNX2 24764292 ChIP−Seq MC3T3 Mouse
MYC 27129775 Chip−Seq CORNEA Mouse
EBNA2 21746931 ChIP−Seq IB4−LCL Human
ATF3 23680149 ChIP−Seq GBM1−GSC Human
SMAD1 26771354 ChIP−Seq mESC Mouse Stem
RELA 24523406 ChIP−Seq FIBROSARCOMA Human
MYBL2 22936984 ChIP−ChIP MESCs Mouse
SRY 25088423 ChIP−ChIP EMBRYONIC GONADS Mouse
SMRT 22465074 ChIP−Seq MACROPHAGES Human
KDM2B 26808549 Chip−Seq SUP−B15 Human
CEBPD 21427703 ChIP−Seq 3T3−L1 Mouse
Nerf2 26677805 Chip−Seq MACROPHAGESS Mouse
PPARG 20887899 ChIP−Seq 3T3−L1 Mouse
E2F1 26619117 ChIP−Seq Hepatocytes Mouse Liver
MECOM 23826213 ChIP−Seq KASUMI Mouse
WT1 25993318 ChIP−Seq PODOCYTE Human
ESR1 22446102 ChIP−Seq UTERUS Mouse
E2F4 17652178 ChIP−ChIP JURKAT Human
FOXM1 26456572 ChIP−Seq MCF−7 Human BreastCancer
ZNF217 24962896 ChIP−Seq MCF−7 Human
LXR 22158963 ChIP−Seq LIVER Mouse
PPARA 22158963 ChIP−Seq LIVER Mouse
RXR 22158963 ChIP−Seq LIVER Mouse
NFKB1 27076634 ChIP−Seq BEAS2B Human Lung Inflammation
NR3C1 27076634 ChIP−Seq BEAS2B Human Lung Inflammation
FOXM1 25889361 ChIP−Seq OE33 AND U2OS Human
FOXM1 23109430 ChIP−Seq U2OS Human
SUZ12 20075857 ChIP−Seq MESCs Mouse
D7
D14
D21
D35
Time point
D
HR
SMAD2/3 21741376 ChIP−Seq EPCs Human
CTCF 18555785 Chip−Seq ESCs Mouse
EZH2 27294783 Chip−Seq ESCs Mouse
EED 16625203 ChIP−ChIP MESCs Mouse
SMAD1 18555785 Chip−Seq ESCs Mouse
RELA 24523406 ChIP−Seq FIBROSARCOMA Human
MYCN 21190229 ChIP−Seq SHEP−21N Human
SUZ12 16625203 ChIP−ChIP MESCs Mouse
MYBL2 22936984 ChIP−ChIP MESCs Mouse
E2F7 22180533 ChIP−Seq HELA Human
AR 21909140 ChIP−Seq LNCAP Human
E2F1 26619117 ChIP−Seq Hepatocytes Mouse Liver
IRF8 27001747 Chip−Seq BMDM Mouse
E2F4 17652178 ChIP−ChIP JURKAT Human
FOXM1 26456572 ChIP−Seq MCF−7 Human BreastCancer
SUZ12 18692474 ChIP−Seq MEFs Mouse
BACH1 22875853 ChIP−PCR HELA AND SCP4 Human
FOXM1 25889361 ChIP−Seq OE33 AND U2OS Human
SUZ12 18692474 ChIP−Seq MESCs Mouse
SUZ12 18974828 ChIP−Seq MESCs Mouse
FOXM1 23109430 ChIP−Seq U2OS Human
SUZ12 20075857 ChIP−Seq MESCs Mouse
−log10(adj. p)
50
40
30
20
10
D7
D14
D21
D35
Time point

Figure S10

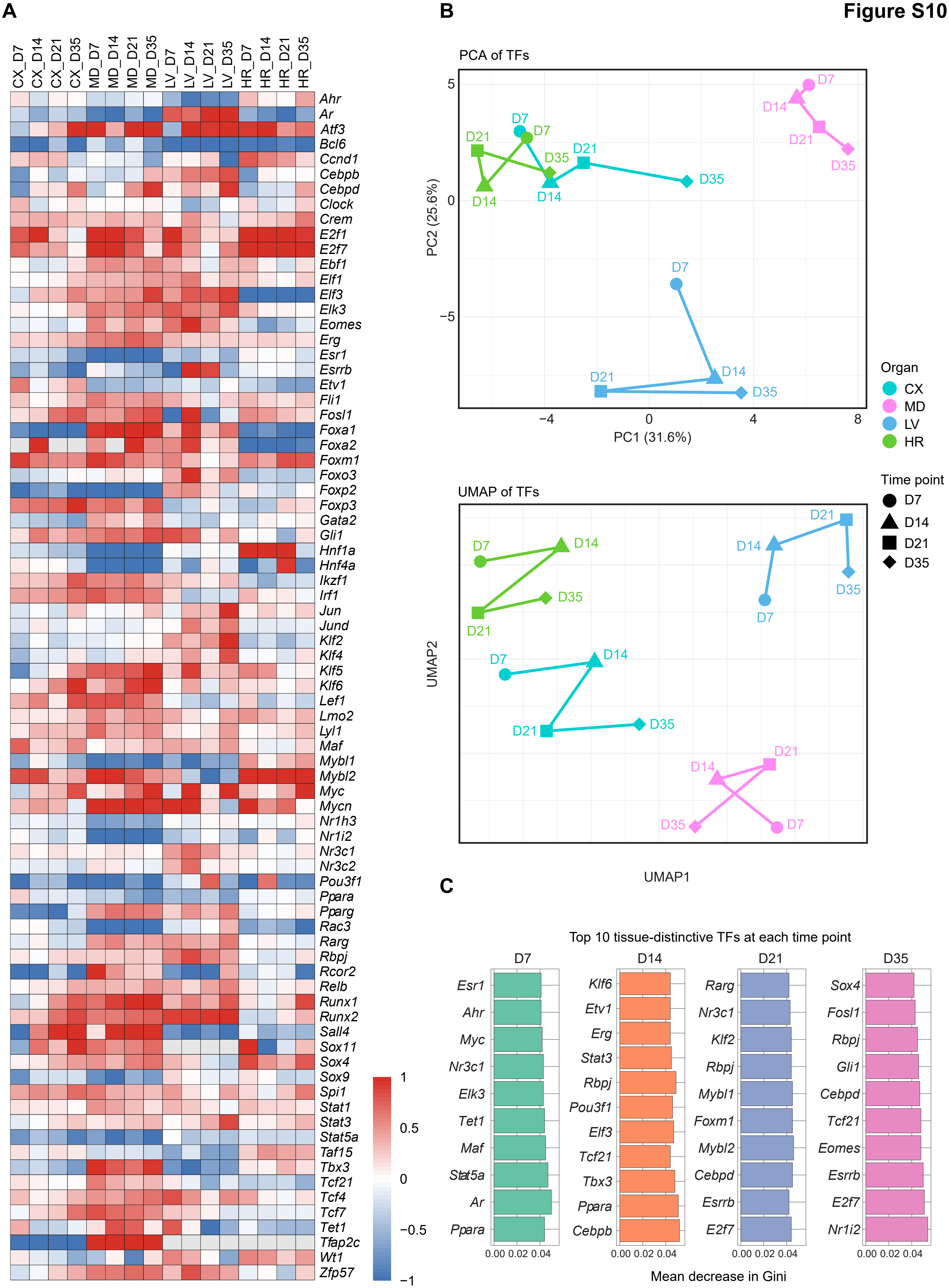

A
CX_D7
CX_D14
CX_D21
CX_D35
MD_D7
MD_D14
MD_D21
MD_D35
LV_D7
LV_D14
LV_D21
LV_D35
HR_D7
HR_D14
HR_D21
HR_D35
Ahr
Ar
Atf3
Bcl6
Ccnd1
Cebpb
Cebpd
Clock
Crem
E2f1
E2f7
Ebf1
Elf1
Elf3
Elk3
Eomes
Erg
Esr1
Esrrb
Etv1
Fli1
Fosl1
Foxa1
Foxa2
Foxm1
Foxo3
Foxp2
Foxp3
Gata2
Gli1
Hnf1a
Hnf4a
Ikzf1
Irf1
Jun
Jund
Klf2
Klf4
Klf5
Klf6
Lef1
Lmo2
Lyl1
Maf
Mybl1
Mybl2
Myc
Mycn
Nr1h3
Nr1i2
Nr3c1
Nr3c2
Pou3f1
Ppara
Pparg
Rac3
Rarg
Rbpj
Rcor2
Relb
Runx1
Runx2
Sall4
Sox11
Sox4
Sox9
Spi1
Stat1
Stat3
Stat5a
Taf15
Tbx3
Tcf21
Tcf4
Tcf7
Tet1
Tfap2c
Wt1
Zfp57
1
0.5
0
−0.5
−1
B
PCA of TFs
PC2 (25.6%)
PC1 (31.6%)
5
0
−5
−4
0
4
8
D7
D14
D21
D35
UMAP of TFs
UMAP2
UMAP1
Organ
CX
MD
LV
HR
Time point
D7
D14
D21
D35
C
Top 10 tissue-distinctive TFs at each time point
D7
Esr1
Ahr
Myc
Nr3c1
Elk3
Tet1
Maf
Stat5a
Ar
Ppara
D14
Klf6
Etv1
Erg
Stat3
Rbpj
Pou3f1
Elf3
Tcf21
Tbx3
Ppara
Cebpb
D21
Rarg
Nr3c1
Klf2
Rbpj
Mybl1
Foxm1
Mybl2
Cebpd
Esrrb
E2f7
D35
Sox4
Fosl1
Rbpj
Gli1
Cebpd
Tcf21
Eomes
Esrrb
E2f7
Nr1i2
0.00 0.02 0.04
Mean decrease in Gini

**Figure S11**

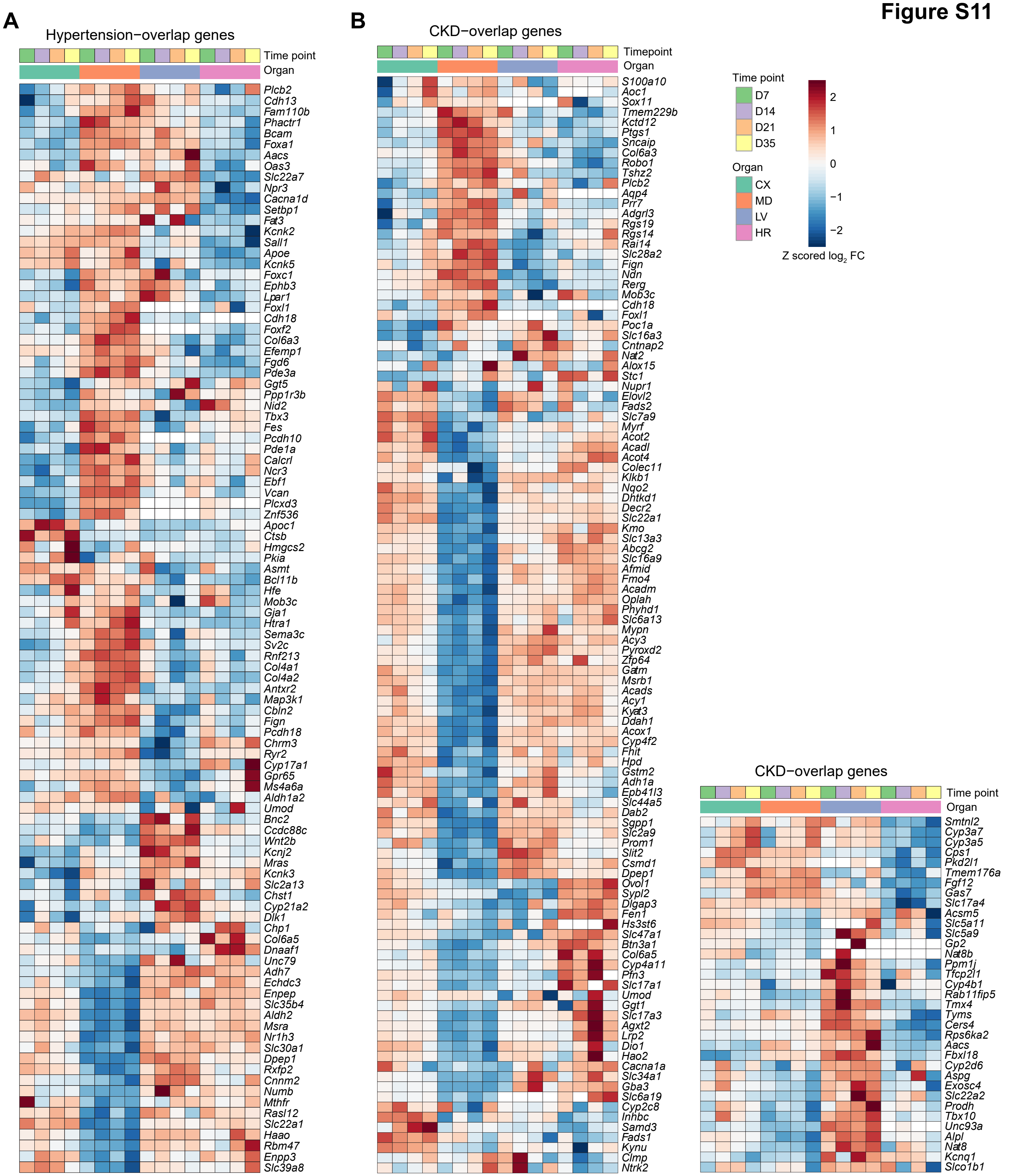
A
Hypertension−overlap genes
Time point
Organ
Plcb2
Cdh13
Fam110b
Phactr1
Bcam
Foxa1
Aacs
Oas3
Slc22a7
Npr3
Cacna1d
Setbp1
Fat3
Kcnk2
Sall1
Apoe
Kcnk5
Foxc1
Ephb3
Lpar1
Foxl1
Cdh18
Foxf2
Col6a3
Efemp1
Fgd6
Pde3a
Ggt5
Ppp1r3b
Nid2
Tbx3
Fes
Pcdh10
Pde1a
Calcrl
Ncr3
Ebf1
Vcan
Plcxd3
Znf536
Apoc1
Ctsb
Hmgcs2
Pkia
Asmt
Bcl11b
Hfe
Mob3c
Gja1
Htra1
Sema3c
Sv2c
Rnf213
Col4a1
Col4a2
Antxr2
Map3k1
Cbln2
Fign
Pcdh18
Chrm3
Ryr2
Cyp17a1
Gpr65
Ms4a6a
Aldh1a2
Umod
Bnc2
Ccdc88c
Wnt2b
Kcnj2
Mras
Kcnk3
Slc2a13
Chst1
Cyp21a2
Dlk1
Chp1
Col6a5
Dnaaf1
Unc79
Adh7
Echdc3
Enpep
Slc35b4
Aldh2
Msra
Nr1h3
Slc30a1
Dpep1
Rxfp2
Cnnm2
Numb
Mthfr
Rasl12
Slc22a1
Haao
Rbm47
Enpp3
Slc39a8
B
CKD−overlap genes
Timepoint
Organ
S100a10
Aoc1
Sox11
Tmem229b
Kctd12
Ptgs1
Sncaip
Col6a3
Robo1
Tshz2
Plcb2
Aqp4
Prr7
Adgrl3
Rgs19
Rgs14
Rai14
Slc28a2
Fign
Ndn
Rerg
Mob3c
Cdh18
Foxl1
Poc1a
Slc16a3
Cntnap2
Nat2
Alox15
Stc1
Nupr1
Elovl2
Fads2
Slc7a9
Myrf
Acot2
Acadl
Acot4
Colec11
Klkb1
Nqo2
Dhtkd1
Decr2
Slc22a1
Kmo
Slc13a3
Abcg2
Slc16a9
Afmid
Fmo4
Acadm
Oplah
Phyhd1
Slc6a13
Mypn
Acy3
Pyroxd2
Zfp64
Gatm
Msrb1
Acads
Acy1
Kyat3
Ddah1
Acox1
Cyp4f2
Fhit
Hpd
Gstm2
Adh1a
Epb41l3
Slc44a5
Dab2
Sgpp1
Slc2a9
Prom1
Slit2
Csmd1
Dpep1
Ovol1
Sypl2
Dlgap3
Fen1
Hs3st6
Slc47a1
Btn3a1
Col6a5
Cyp4a11
Pfn3
Slc17a1
Umod
Ggt1
Slc17a3
Agxt2
Lrp2
Dio1
Hao2
Cacna1a
Slc34a1
Gba3
Slc6a19
Cyp2c8
Inhbc
Samd3
Fads1
Kynu
Clmp
Ntrk2
Time point
D7
D14
D21
D35
Organ
CX
MD
LV
HR
2
1
0
−1
−2
Z scored log2 FC
CKD−overlap genes
Time point
Organ
Smtnl2
Cyp3a7
Cyp3a5
Cps1
Pkd2l1
Tmem176a
Fgf12
Gas7
Slc17a4
Acsm5
Slc5a11
Slc5a9
Gp2
Nat8b
Ppm1j
Tfcp2l1
Cyp4b1
Rab11fip5
Tmx4
Tyms
Cers4
Rps6ka2
Aacs
Fbxl18
Cyp2d6
Aspg
Exosc4
Slc22a2
Prodh
Tbx10
Unc93a
Alpl
Nat8
Kcnq1
Slco1b1

**Figure S12**

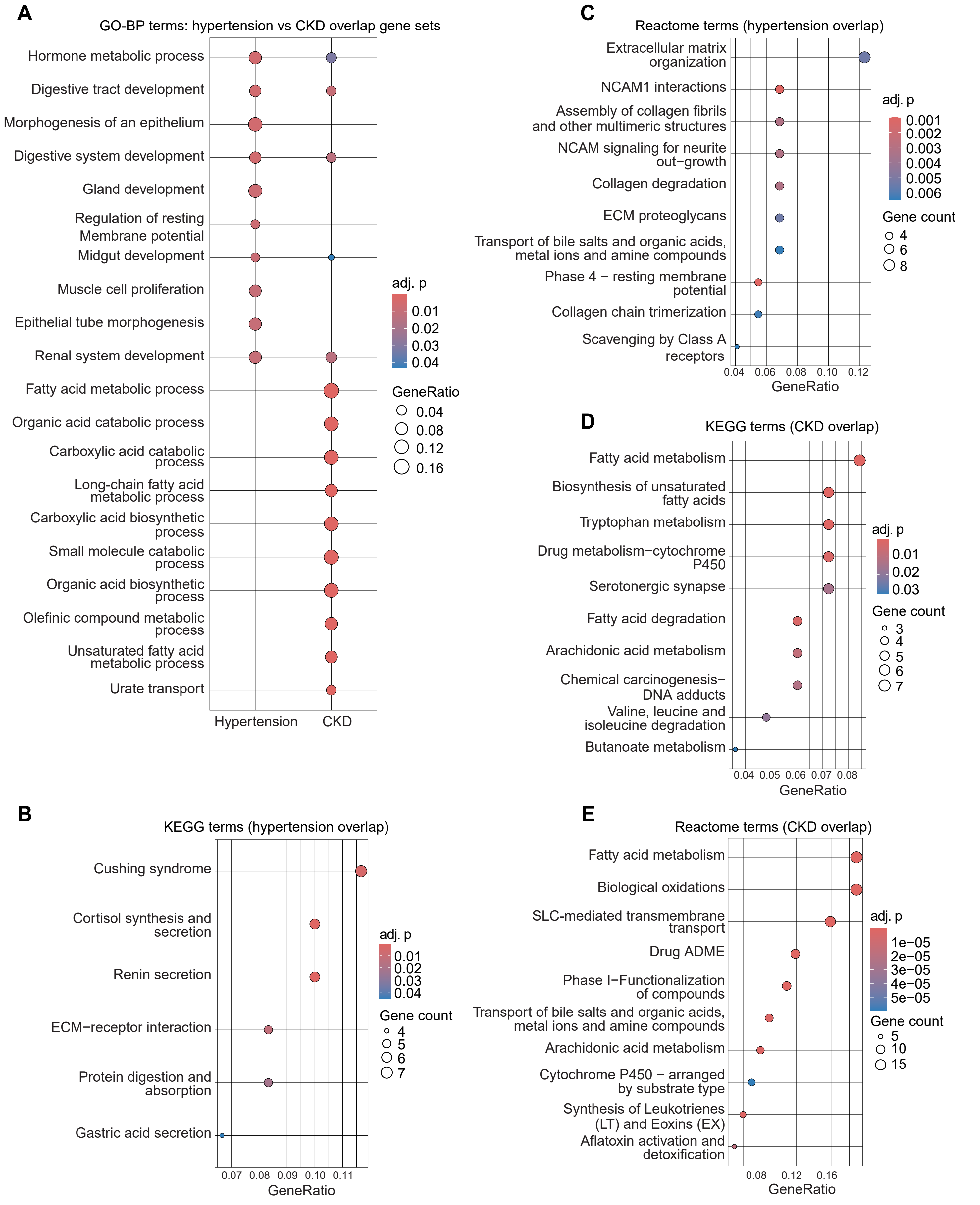

A
GO-BP terms: hypertension vs CKD overlap gene sets
Hormone metabolic process
Digestive tract development
Morphogenesis of an epithelium
Digestive system development
Gland development
Regulation of resting Membrane potential
Midgut development
Muscle cell proliferation
Epithelial tube morphogenesis
Renal system development
Fatty acid metabolic process
Organic acid catabolic process
Carboxylic acid catabolic process
Long-chain fatty acid metabolic process
Carboxylic acid biosynthetic process
Small molecule catabolic process
Organic acid biosynthetic process
Olefinic compound metabolic process
Unsaturated fatty acid metabolic process
Urate transport
Hypertension
CKD
adj. p
0.01
0.02
0.03
0.04
GeneRatio
0.04
0.08
0.12
0.16
B
KEGG terms (hypertension overlap)
Cushing syndrome
Cortisol synthesis and secretion
Renin secretion
ECM−receptor interaction
Protein digestion and absorption
Gastric acid secretion
0.07 0.08 0.09 0.10 0.11
GeneRatio
adj. p
0.01
0.02
0.03
0.04
Gene count
4
5
6
7
C
Reactome terms (hypertension overlap)
Extracellular matrix organization
NCAM1 interactions
Assembly of collagen fibrils and other multimeric structures
NCAM signaling for neurite out−growth
Collagen degradation
ECM proteoglycans
Transport of bile salts and organic acids, metal ions and amine compounds
Phase 4 − resting membrane potential
Collagen chain trimerization
Scavenging by Class A receptors
0.04 0.06 0.08 0.10 0.12
GeneRatio
adj. p
0.001
0.002
0.003
0.004
0.005
0.006
Gene count
4
6
8
D
KEGG terms (CKD overlap)
Fatty acid metabolism
Biosynthesis of unsaturated fatty acids
Tryptophan metabolism
Drug metabolism−cytochrome P450
Serotonergic synapse
Fatty acid degradation
Arachidonic acid metabolism
Chemical carcinogenesis− DNA adducts
Valine, leucine and isoleucine degradation
Butanoate metabolism
0.04 0.05 0.06 0.07 0.08
GeneRatio
adj. p
0.01
0.02
0.03
Gene count
3
4
5
6
7
E
Reactome terms (CKD overlap)
Fatty acid metabolism
Biological oxidations
SLC-mediated transmembrane transport
Drug ADME
Phase I−Functionalization of compounds
Transport of bile salts and organic acids, metal ions and amine compounds
Arachidonic acid metabolism
Cytochrome P450 − arranged by substrate type
Synthesis of Leukotrienes (LT) and Eoxins (EX)
Aflatoxin activation and detoxification
0.08 0.12 0.16
GeneRatio
adj. p
1e−05
2e−05
3e−05
4e−05
5e−05
Gene count
5
10
15

**Figure S13**

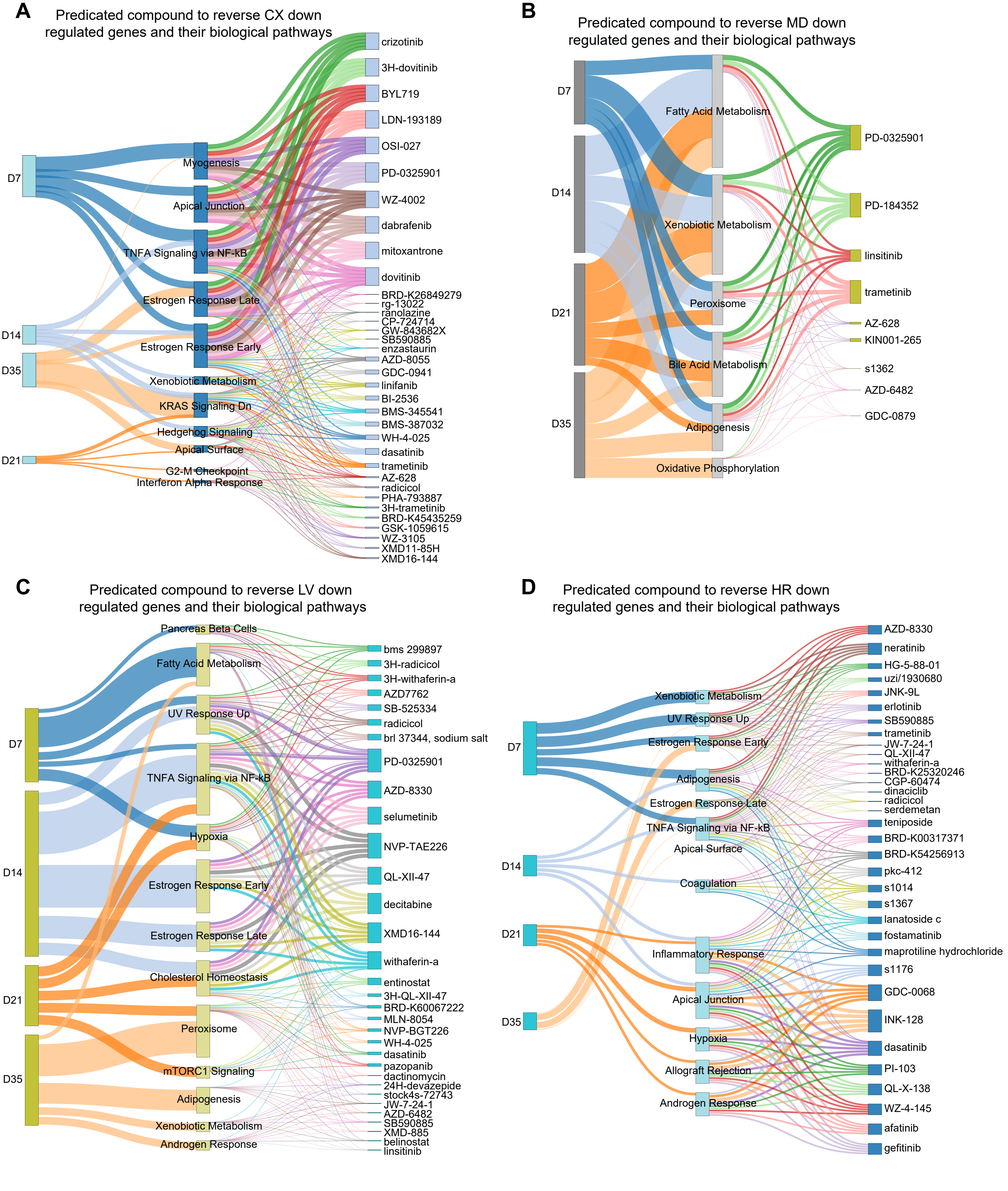

A
Predicated compound to reverse CX down regulated genes and their biological pathways
D7
D14
D35
D21
Myogenesis
Apical Junction
TNFA Signaling via NF-kB
Estrogen Response Late
Estrogen Response Early
Xenobiotic Metabolism
KRAS Signaling Dn
Hedgehog Signaling
Apical Surface
G2-M Checkpoint
Interferon Alpha Response
crizotinib
3H-dovitinib
BYL719
LDN-193189
OSI-027
PD-0325901
WZ-4002
dabrafenib
mitoxantrone
dovitinib
BRD-K26849279
rg-13022
ranolazine
CP-724714
GW-843682X
SB590885
enzastaurin
AZD-8055
GDC-0941
linifanib
BI-2536
BMS-345541
BMS-387032
WH-4-025
dasatinib
trametinib
AZ-628
radicicol
PHA-793887
3H-trametinib
BRD-K45435259
GSK-1059615
WZ-3105
XMD11-85H
XMD16-144
B
Predicated compound to reverse MD down regulated genes and their biological pathways
D7
D14
D21
D35
Fatty Acid Metabolism
Xenobiotic Metabolism
Peroxisome
Bile Acid Metabolism
Adipogenesis
Oxidative Phosphorylation
PD-0325901
PD-184352
linsitinib
trametinib
AZ-628
KIN001-265
s1362
AZD-6482
GDC-0879
C
Predicated compound to reverse LV down regulated genes and their biological pathways
D7
D14
D21
D35
Pancreas Beta Cells
Fatty Acid Metabolism
UV Response Up
TNFA Signaling via NF-kB
Hypoxia
Estrogen Response Early
Estrogen Response Late
Cholesterol Homeostasis
Peroxisome
mTORC1 Signaling
Adipogenesis
Xenobiotic Metabolism
Androgen Response
bms 299897
3H-radicicol
3H-withaferin-a
AZD7762
SB-525334
radicicol
brl 37344, sodium salt
PD-0325901
AZD-8330
selumetinib
NVP-TAE226
QL-XII-47
decitabine
XMD16-144
withaferin-a
entinostat
3H-QL-XII-47
BRD-K60067222
MLN-8054
NVP-BGT226
WH-4-025
dasatinib
pazopanib
dactinomycin
24H-devazepide
stock4s-72743
JW-7-24-1
AZD-6482
SB590885
XMD-885
belinostat
linsitinib
D
Predicated compound to reverse HR down regulated genes and their biological pathways
D7
D14
D21
D35
Xenobiotic Metabolism
UV Response Up
Estrogen Response Early
Adipogenesis
Estrogen Response Late
TNFA Signaling via NF-kB
Apical Surface
Coagulation
Inflammatory Response
Apical Junction
Hypoxia
Allograft Rejection
Androgen Response
AZD-8330
neratinib
HG-5-88-01
uzi/1930680
JNK-9L
erlotinib
SB590885
trametinib
JW-7-24-1
QL-XII-47
withaferin-a
BRD-K25320246
CGP-60474
dinaciclib
radicicol
serdemetan
teniposide
BRD-K00317371
BRD-K54256913
pkc-412
s1014
s1367
lanatoside c
fostamatinib
maprotiline hydrochloride
s1176
GDC-0068
INK-128
dasatinib
PI-103
QL-X-138
WZ-4-145
afatinib
gefitinib
